# ATP-responsive biomolecular condensates tune bacterial kinase signaling

**DOI:** 10.1101/2020.08.09.232405

**Authors:** Saumya Saurabh, Trisha N. Chong, Camille Bayas, Peter D. Dahlberg, Heather N. Cartwright, W. E. Moerner, Lucy Shapiro

## Abstract

Biomolecular condensates formed via phase separation enable spatial and temporal organization of enzyme activity. The emergent properties of many condensates have been shown to be responsive to intracellular Adenosine triphosphate (ATP) levels, although the consequences of such mechanisms on enzyme activity are unknown. Here, we show that ATP depletion promotes phase separation in condensates composed of a disordered protein, thereby enhancing the activity of a client kinase enabling robust signaling and maintenance of viability under the stress posed by nutrient scarcity. We propose that a diverse repertoire of condensates can serve as control knobs to tune multivalency and reactivity in response to the metabolic state of bacterial cells.

**One Sentence Summary:** Bacterial condensates boost kinase activity under ATP depletion

## Main Text

All cells translate metabolic cues into molecular signals for successful proliferation in dynamic environments. Biomolecular condensates formed by phase separation of multivalent disordered proteins enable cells to regulate biochemical signals in response to their environment (*1, 2*). A subset of disordered proteins exhibit an ATP-dependent multivalency (*3*) that is independent of ATP’s role as a substrate for active processes. Yet, it remains unknown whether modulation of multivalency by ATP has a functional relevance for ATP-consuming enzymes compartmentalized within condensates. Bacteria that survive over a broad range of conditions and serve as simple yet powerful systems to test the role of ATP-depletion on the activity of compartmentalized ATP-consuming enzymes. Here, we demonstrate that the activity of a bacterial ATP-dependent kinase, responds to the emergent ATP-dependent multivalency of the condensate that it inhabits. ATP depletion promotes phase separation, enforces kinase compartmentalization, and sustains kinase activity under nutrient scarcity. Therefore, co-option of the dual roles of ATP renders cellular fitness under stress induced by nutrient depletion.

The free-living bacterium *Caulobacter crescentus* (hereafter, *Caulobacter*) thrives in aquatic habitats and divides asymmetrically yielding a sessile, stalked cell and a motile, swarmer cell during each cell cycle (Fig. 1A). The swarmer cell is incapable of DNA replication and cell division until it differentiates into a stalked cell (*4*). Differentiation is marked by the loss of flagellum and initiation of stalk biogenesis at the pole. Concomitantly, the polar microdomain composed of the disordered protein PopZ exhibits a change in its client protein composition (*5*). A newly expressed protein, SpmX, localizes to the stalk-bearing pole through a direct interaction with PopZ (*6, 7*). SpmX is an integral membrane disordered protein that subsequently localizes a membrane associated histidine kinase DivJ to the stalked pole. The autokinase activity of DivJ is critical for cell division and depends on its stable localization to the stalked pole, thereby establishing SpmX as a key regulator of the stalked cell developmental program (*6*). Cells harboring a deletion of SpmX, exhibit slow growth rates and aberrant cell division, phenocopying a DivJ mutant (*6*). Despite the phenotypic relevance, the precise mechanisms of SpmX-dependent DivJ localization and activity are poorly understood.

**Fig. 1.**
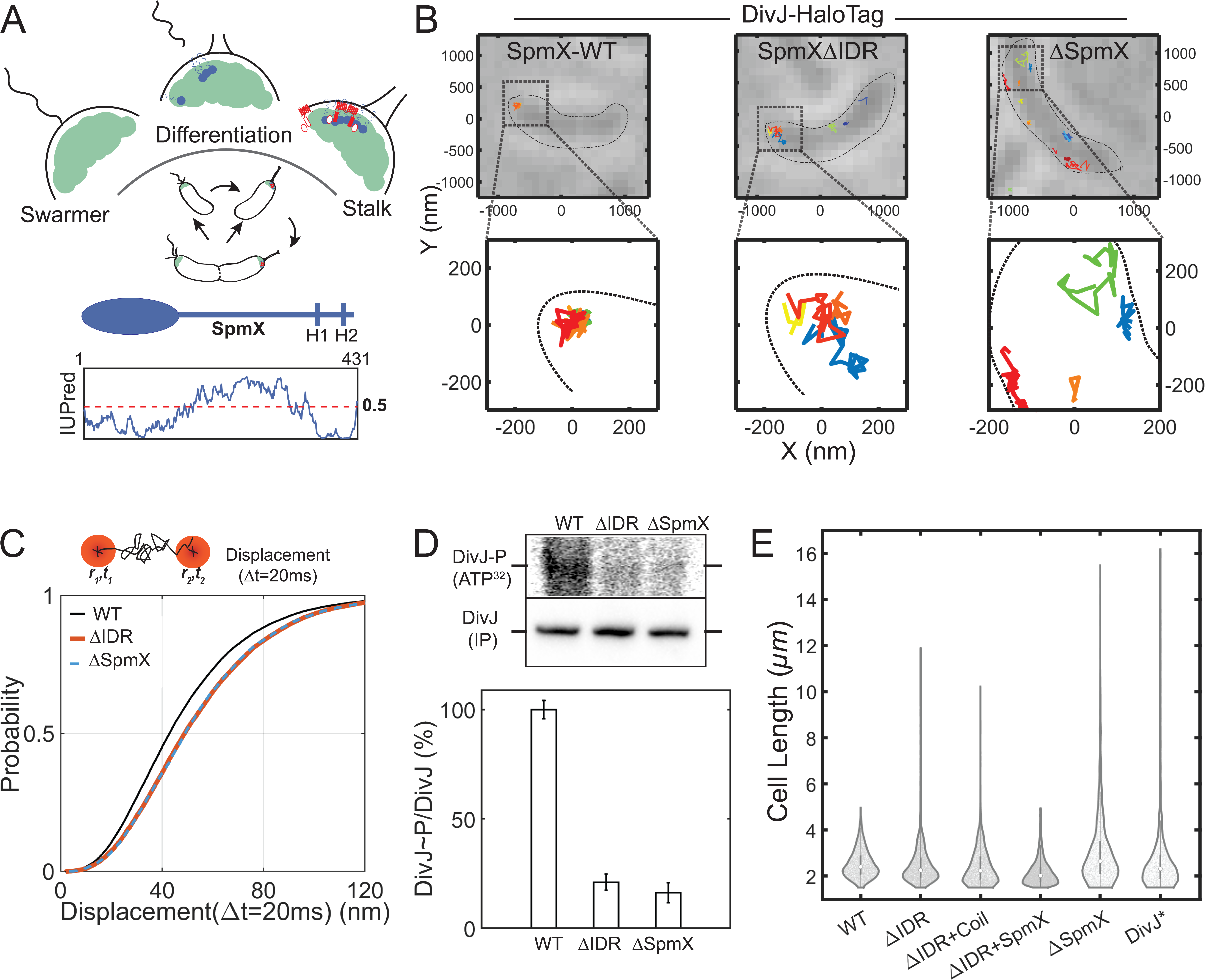
SpmX-IDR sequesters and stimulates DivJ kinase at the pole. (**A**) A schematic of the *Caulobacter* cell cycle showing the key steps of cell division. Zoomed in schematic above shows the pole during differentiation from swarmer to stalked cell. PopZ (green), SpmX (blue), and the DivJ kinase (red). Shown below, is the domain organization and disorder prediction for SpmX calculated using IUPred. Transmembrane helices are denoted as H1 and H2. Asymmetric localization of the disordered protein SpmX and kinase DivJ at the stalk-bearing pole is critical for cell cycle progression. (**B**) (Top) Representative white light images of live *Caulobacter* cells expressing an endogenous DivJ-HaloTag fusion in strains bearing different SpmX perturbations. Representative single DivJ trajectories are overlaid on the white light images. Approximate cell boundary is indicated using black dashed line. (Bottom) a zoomed in view of four polar trajectories (for WT-SpmX and SpmXΔIDR) and non-polar trajectories (for ΔSpmX). Each trajectory is shown as a 2D projection and in a different color. DivJ exhibits tightly localized polar trajectories in WT cells while more diffuse polar trajectories in SpmXΔIDR cells. **(C)** Cumulative distribution function (CDF) of displacements in 20 ms intervals obtained from DivJ trajectories in WT, SpmXΔIDR, and ΔSpmX cells. Inset shows a zoomed in view of the graph highlighting the overlapping CDFs for SpmXΔIDR and ΔSpmX. The number of polar DivJ trajectories analyzed were 974, 443 and 291, for WT, SpmXΔIDR and ΔSpmX cells, respectively. In the absence of SpmX-IDR, DivJ exhibited higher diffusivity in the polar microdomain. **(D)** *In vivo* phosphorylation measurement of DivJ in SpmX perturbations. DivJ phosphorylation levels in the bar plot are expressed as a ratio of DivJ∼P (intensity in the top gel (ATP(γ−^32^P)) to DivJ concentration (immunoprecipitation using an antibody against DivJ-HaloTag) and are normalized to the ratio observed in WT cells. ΔIDR denotes SpmXΔIDR. Error bars denote standard error of the mean from three biological replicates. SpmX-IDR stimulates DivJ activity *in vivo*. **(E)** Distribution of lengths for *Caulobacter* cells grown in M2 minimal media supplemented with 0.2 mM glucose for 4 hours, followed by phase contrast microscopy. The white dot on dark gray bars within violin plots denotes the median value of the distribution. ΔIDR denotes SpmXΔIDR; Coil denotes a strain endogenously expressing the CoilZ fusion of SpmXΔIDR and a CoilY fusion of DivJ; ΔIDR+SpmX denotes the SpmXΔIDR strain with a mild expression of WT SpmX; DivJ* denotes DivJ(H338A) catalytic mutant. A total of 2000-7000 cells were analyzed in each case from three biological replicates. SpmXΔIDR cells mimic a DivJ mutant phenotype under glucose depletion.

How is the activity of relevant kinases such as DivJ regulated under low intracellular ATP concentrations? In addition to its role as a substrate for active cellular processes, ATP has been shown to regulate the solubility of eukaryotic proteins containing disordered domains (*3*). However, it is unclear whether the role of ATP as a solute that regulates multivalency of disordered proteins is conserved in bacteria and if such mechanisms affect ATP-utilizing enzymes sequestered via disordered proteins. The ATP-dependent kinase reaction of DivJ, in the context of its interactions with SpmX and PopZ, provides a system to test the impact of emergent properties of condensates on the activity of an enzyme that uses ATP as a substrate.

Here, we elucidate the molecular basis for DivJ activity under low ATP conditions. We show that DivJ compartmentalization is mediated by a disordered domain in SpmX. SpmX, and its localization determinant, PopZ both contain disordered domains that promote multivalent interactions and condensation through phase separation. The multivalence of SpmX and PopZ are shown to be emergent properties in that they can be tuned by ATP within a physiological range. The interplay between ATP levels and SpmX multivalency enabled robust modulation of DivJ activity under low ATP concentrations. Taken together, our findings underscore the role of ATP-dependent multivalency in disordered proteins and its relevance for cellular signaling as a function of nutrient availability.

## Results

### SpmX-IDR sequesters DivJ to the pole and stimulates DivJ kinase activity

SpmX has been shown to be indispensable for both DivJ polar localization and kinase activity (*6*). In addition to a structured, lysozyme-like domain that interacts with PopZ (*7*), SpmX contains an intrinsically disordered region (IDR) of ∼200 amino acids (Fig. 1A), whose relevance remains unclear. *In vitro*, the affinity of DivJ for SpmX is lowered in the absence of the SpmX-IDR (*7*). To determine whether SpmX-IDR influences the polar localization of DivJ *in vivo*, we performed three-dimensional (3D) single-molecule tracking (*8*) of DivJ (materials and methods), endogenously fused to HaloTag and labeled using Janelia Fluor 549 (JF549-HaloTag) (*9*) in SpmX wild type (WT), SpmXΔIDR, or *spmX* deletion (ΔSpmX) strains (Fig. 1B). The DivJ-HaloTag construct, controlled by the native *divJ* promoter fully complemented a Δ*divJ* strain. DivJ exhibited tightly localized polar trajectories in WT cells but diffuse polar trajectories in the SpmXΔIDR cells (Fig. 1B and fig. S1A), indicative of a faster polar diffusion of DivJ in the absence of the SpmX-IDR. The absence of the SpmX-IDR resulted in a decrease in DivJ’s polar concentration. While 60% of all the DivJ trajectories were observed at the pole in WT cells, this number dropped to 20% in SpmXΔIDR cells, and to 10% in ΔSpmX cells (Fig. S1B). Owing to the small polar diffusivities and limited trajectory lengths, DivJ tracks were analyzed using the cumulative distribution function (CDF) of displacements (*10*) in 20 ms time intervals (Fig. 1C). CDF analyses showed that DivJ displacements in the polar microdomain were larger in both ΔSpmX and SpmXΔIDR cells compared to WT cells (Fig. 1C and fig. S1C-D, supplementary text), indicating that SpmX, and specifically the SpmX-IDR, increase the residence time of DivJ at the pole. Together with previous data, these results establish a modular role of SpmX in which SpmX polar localization is mediated by an interaction between the SpmX lysozyme-like domain and PopZ (*7*), while the interaction between SpmX-IDR and DivJ results in DivJ’s sequestration within the polar microdomain.

Cells lacking SpmX exhibit a dramatic decrease in DivJ phosphorylation, and compromised viability (*6*). To determine if polar sequestration of DivJ via SpmX-IDR regulates DivJ kinase activity, we measured DivJ phosphorylation levels *in vivo* in WT, ΔSpmX, or SpmXΔIDR strains using radiolabeled ATP and immunoprecipitation (Fig. 1D, materials and methods). Compared to WT cells, SpmXΔIDR strains exhibited an 80% reduction in the levels of phosphorylated DivJ, which was comparable to the levels observed in the *spmX* deletion strain (Fig. 1D). The dependence of DivJ kinase activity on the SpmX-IDR could be a result of a direct conformational regulation of its catalytic domains by the SpmX-IDR or DivJ sequestration within the polar microdomain. Therefore, we tested if the autokinase activity of DivJ was enhanced by sequestration in its physiological topology *in vitro*. The cytoplasmic domains of DivJ (DivJ(ΔTM)) were tethered to liposomes via an N-terminal His_6_-Tag (materials and methods) (*11*). In this assay, we observed a density-dependent increase in DivJ kinase activity, suggesting that DivJ sequestration can enhance its kinase activity *in vitro* (Fig. S1E-F). While an allosteric regulation of DivJ activity via SpmX-IDR cannot be ruled out, it is plausible that polar sequestration of DivJ via the SpmX-IDR enhances DivJ kinase activity through the effects of increased local concentration that contribute to cooperativity in substrate binding and turnover.

Cells with disrupted DivJ localization, or inhibited kinase activity via mutations to DivJ are filamentous due to disruption of cell division (*6, 12*). Accordingly, we expected SpmXΔIDR cells to be filamentous and phenocopy a SpmX deletion or a DivJ kinase mutant (Fig. S1G). However, the filamentous cell phenotype in SpmXΔIDR cells was observed only under glucose limitation and in stationary phase. Accordingly, we used the distribution of cell lengths for strains grown in low glucose media as a robust phenotypic readout to elucidate the role of SpmX-IDR in DivJ regulation *in vivo*. While less than 1% WT cells were filamentous (> 4 µm), ΔSpmX, SpmXΔIDR, and DivJ kinase mutants exhibited an increased number of aberrantly dividing cells under low glucose (Fig. 1E). The filamentous cell phenotype could be rescued by low expression of WT SpmX in SpmXΔIDR strains (Fig. 1E and fig. S1I). To distinguish between the effect of SpmX-IDR mediated polar sequestration from other regulatory mechanisms, we constructed a strain where DivJ polar recruitment was independent of the SpmX-IDR. We utilized cognate coiled domains with strong binding affinity for this purpose (*13*). A strain with SpmXΔIDR fused to CoilY and DivJ fused to CoilZ exhibited WT-like distribution of DivJ polar localization (materials and methods, Fig. S1H). However, DivJ localization via a structured coil domain could not rescue the filamentous cell phenotype (Fig. 1E), suggesting that SpmX-IDR sequesters and regulates DivJ in response to glucose depletion.

Taken together, the data show that SpmX-IDR is critical for sequestering DivJ at the pole, thereby enhancing its cooperativity and kinase activity. Additionally, the SpmX-IDR mediated cell division phenotype could not be rescued by artificial polar sequestration of DivJ via structured domains, suggesting an allosteric regulatory mechanism that is critical under glucose depletion. The intriguing observation of glucose dependent regulation of DivJ activity necessitated an investigation of SpmX and PopZ IDR properties that could be related to glucose metabolism.

### SpmX and PopZ form interacting polar condensates via multivalent interactions

The PopZ microdomain has been shown to selectively sequester client proteins at the cell poles (*14*). SpmX is one such client protein that localizes to the stalked pole during differentiation. Both SpmX and PopZ have IDR sequences rich in proline and acidic residues (Fig. 2A). Multiple IDR sequences have been shown to exhibit multivalent interactions that enable formation of biomolecular condensates via phase separation and gelation (*15, 16*). Accordingly, we determined if SpmX and PopZ could form condensates under physiological conditions *in vitro*. Purified PopZ and SpmX exhibited a concentration dependent increase in turbidity, suggesting the presence of protein assemblies (Fig. S2A, materials and methods). Microscopic observation of sparsely labeled proteins showed that while DivJ remained soluble in a physiological buffer, SpmX, and PopZ formed spherical condensates under similar conditions (Fig. 2B, materials and methods). Microscopic observation of purified protein domains revealed that the IDR of SpmX (Amino acids (AA) 156-355) and PopZ (AA 1-102) were necessary and sufficient for condensate formation *in vitro* (Fig. S2B). Sparsely labeled DivJ partitioned into and was enriched in condensates formed via SpmX or eYFP-fused to SpmX-IDR (Fig. 2C, Videos S1-S2, Supplementary text), suggesting that SpmX-IDR can directly interact with DivJ *in vitro*, supporting the observation of SpmX-IDR dependent DivJ diffusion *in vivo* (Fig. 1B and C).

**Fig. 2.**
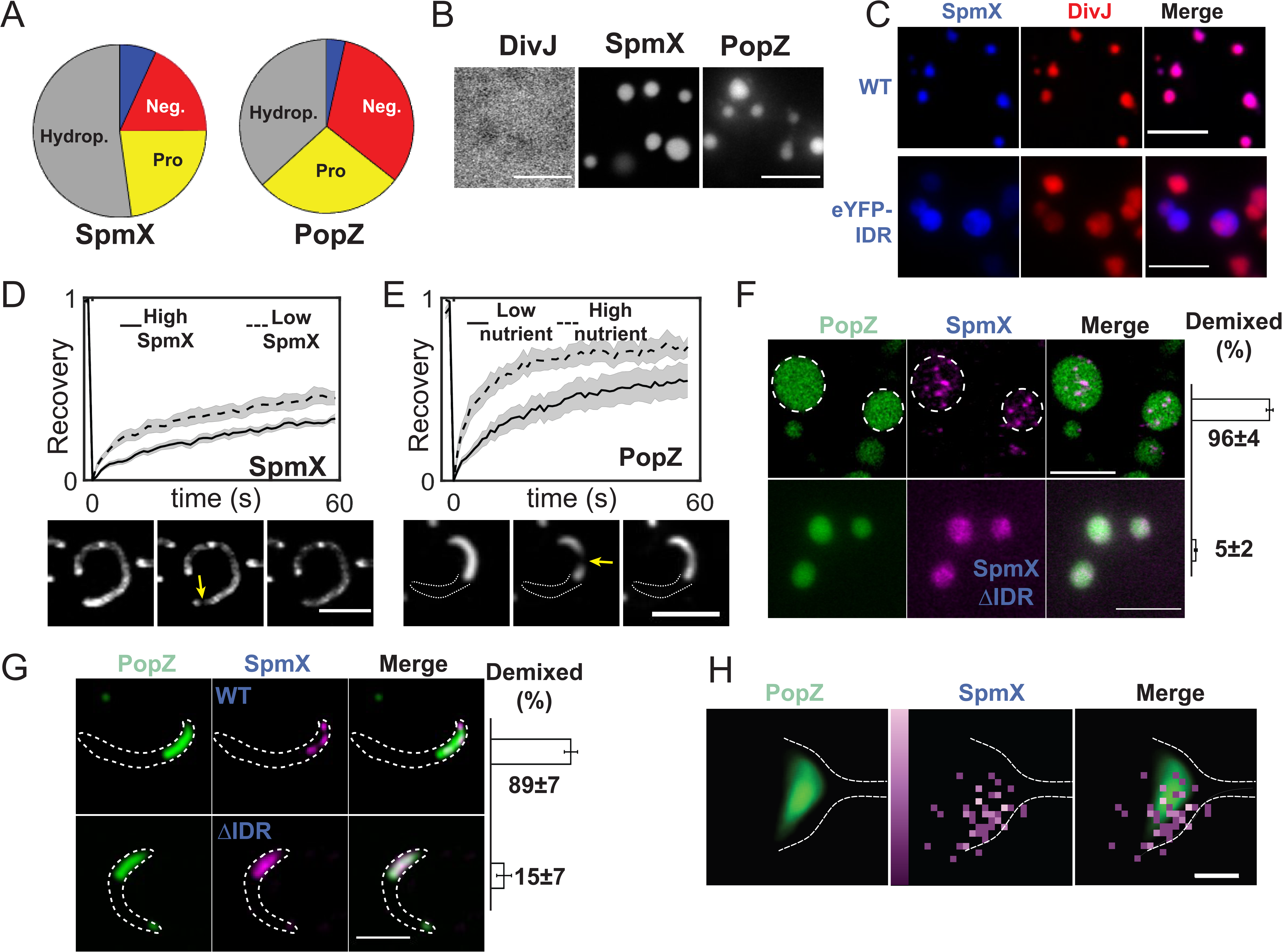
SpmX and PopZ form coexisting yet demixed condensates via multivalent interactions. (All scale bars unless noted, are 5 µm) **(A)** Pie charts depicting the percentage of Hydrophobic (Hydrop., gray), Prolines (Pro., yellow), Negatively charged (Neg., red) and positively charged (blue) residues in SpmX-IDR (AA 156-355) and PopZ-IDR (AA 1-102). **(B)** Representative fluorescence images of purified 5 µM DivJ (ΔTM, 1% labeled with Atto488), 5 µM SpmX (ΔTM, 5% labeled with Cy3), and 5 µM PopZ (1% labeled with Atto488, heated at 95 °C for 8 mins) reconstituted in a physiological buffer (50 mM HEPES-KOH, pH 7.4, 0.1 M KCl). DivJ exhibited diffuse signal while SpmX and PopZ formed round condensates. **(C)** Fluorescence micrographs showing 5 µM DivJ (ΔTM, 1% labeled with Atto488) incubated with 5 µM SpmX (ΔTM, 5% labeled with Cy3) (top) or 5 µM DivJ (ΔTM, 1% labeled with Cy3) incubated with 5 µM eYFP-tagged SpmX-IDR (bottom) in a physiological buffer for 30 mins. DivJ is enriched within SpmX and SpmX-IDR condensates. **(D)** Internal rearrangement of eYFP-labeled SpmX clusters in live *Caulobacter* cells harboring a *popZ* deletion assayed by fluorescence recovery after photobleaching (FRAP). Fluorescence micrographs before and after bleaching are shown below the plot. Yellow arrow points to bleached spot in the first frame after photobleaching. Analysis of recovery profiles are shown for low induction of SpmX (dashed line, N = 11 cells) and high induction of SpmX (solid lines, N = 12 cells). Low and high induction conditions were achieved by supplementing minimal media with 0.03% or 0.3% Xylose, respectively. Cells were induced under these conditions for 4 hours. Gray shadow around the recovery curves represents the standard error of the mean. SpmX clusters exhibited concentration dependent internal dynamics indicative of tunable multivalency *in vivo*. **(E)** Internal rearrangement of eYFP-labeled PopZ condensates in live *Caulobacter* cells assayed by FRAP under two rich and minimal media. Fluorescence micrographs before and after bleaching are shown. Yellow arrow points to bleached spot, while white dashed line denotes an approximate cell boundary. Analysis of recovery profiles are shown for rich media (blue, N = 11 cells) and minimal media (black, N = 11 cells). Gray shadow around the recovery curves represents the standard error of the mean. PopZ exhibits a nutrient dependent internal rearrangement. **(F)** Representative super-resolution images of 5 µM PopZ (1% Atto-488 labeled, heated at 95 °C for 8 mins) incubated with 500 nM (top) SpmX (ΔTM, 5% labeled with Cy3) or (bottom) SpmXΔIDR (ΔTM, 5% labeled with Cy3) proteins in a physiological buffer for 30 mins. Bar plot on the right indicates the % of PopZ condensates that exhibited more than one SpmX cluster within them, serving as a proxy for demixing. For each sample, a total of ∼400 condensates were analyzed. Error bars denote the standard deviation calculated from three biological replicates. PopZ-SpmX coacervates exhibited demixed SpmX clusters while SpmXΔIDR remained uniformly distributed within PopZ condensates *in vitro*. **(G)** Super-resolution images of cells expressing mCherry-PopZ on a high copy plasmid and endogenously tagged SpmX-dL5 (top) or SpmXΔIDR-dL5 (bottom) in a *spmX* deletion strain. WT SpmX formed multiple clusters associated with the PopZ microdomain, while SpmXΔIDR remained uniformly distributed within PopZ microdomain. Bar plot on the right indicates the % of cells with PopZ microdomain larger than 1 µm that exhibited more than one SpmX cluster within them, serving as a proxy for demixing. For each sample, a total of ∼230 cells were analyzed. Error bars denote the standard deviation calculated from three biological replicates. SpmX clusters remain demixed within the PopZ microdomain. **(H)** (left) Manually annotated ribosome excluded region serving as a proxy for the PopZ microdomain from cryogenic electron tomography of four cells. White dashed line denotes an approximate cell boundary. (middle) Localizations of SpmX-PAmKate fusion from correlative cryogenic single molecule imaging pooled from the same four aligned stalked cells. Each pixel is ∼14 nm. Color bar on the left of SpmX image denotes the number of localizations binned into each pixel (range 1 to 3 localizations from dark to light shade). (right) In the overlaid image, SpmX localizations exhibit a non-uniform distribution on one side of the PopZ microdomain suggestive of demixing. (Scale bar 100 nm).

*In vitro*, SpmX condensates exhibited spherical morphologies in solution (viewed away from the glass) and relaxed on the aminosilane treated glass surface. SpmX condensates exhibited moderate internal dynamics just after relaxing on the glass, and subsequently underwent rapid gelation, as measured via fluorescence recovery after photobleaching (FRAP) (Fig. S2C, materials and methods). Rapid fusion events between SpmX condensates were observed within the first minute of condensates fusing on glass while condensate growth by ripening was observed over longer time scales (Fig. S2D). These data suggest that SpmX self-association is enhanced after fusing to glass, resulting in an increase in SpmX concentration within the condensate over time. *In vivo*, we found that over-expression of SpmX-eYFP in cells lacking PopZ led to the formation of SpmX clusters in a concentration dependent manner (Fig. S2E). In accord with *in vitro* measurements, SpmX clusters exhibited concentration dependent internal rearrangements *in vivo*, suggestive of a context dependent multivalency (Fig. 2D). Notably, fluorescent clusters were not observed upon over-expressing the trans-membrane (TM) helices of SpmX fused to eYFP, implicating the cytoplasmic domain in cluster formation (Fig. S2F). *In vitro*, SpmX exhibited a protein and salt concentration dependent phase behavior. At protein concentrations above 5 µM (0.1 M-0.5 M KCl), condensates deviated from the spherical shape and wetted the glass surface more, marking a region of instability on the phase diagram or gelation suppressing phase separation (*2*) (Fig. S2G).

Unlike SpmX, PopZ condensates reconstituted directly from purified stocks deviated from a spherical morphology *in vitro* (Fig. S2H, 23 °C sample). 3D confocal imaging revealed the presence of dynamically arrested condensates (Fig. S2H, Videos S3-S4), reminiscent of poly-Arginine-RNA assemblies that could be relaxed using thermal energy (*17*). Heating condensates that exhibited a “beads on a string” morphology at 95 °C for 8 mins led to the formation of round condensates (Fig. 2B, Fig. S2H, materials and methods), suggestive of a lower critical solution temperature behavior (*18*), whereby higher temperature promotes multivalent protein interactions that result in a collapsed, spherical state. The two forms of PopZ exhibited distinct internal rearrangements and time dependent behavior (Fig. S2I-J, supplementary text). Over-expression of mCherry-PopZ in *Caulobacter* cells led to an increase in the size of the PopZ microdomain extending from the pole. This observation is in line with the space filling nature of PopZ (*8*), and allowed the measurement of internal dynamics within PopZ microdomain *in vivo*. Under the conditions tested, PopZ did not exhibit a concentration dependent diffusivity. However, for comparable levels of protein expression, we observed a significant difference in PopZ internal dynamics between cells transferred to high versus low glucose containing media post protein expression, suggesting a role of metabolites in PopZ diffusion (Fig. 2E). *In vitro*, PopZ condensates exhibited a salt and concentration dependent behavior (Fig. S2K). Taken together, the data show that both SpmX, and its localization determinant, PopZ utilize multivalent interactions from their IDRs to form condensates that are sensitive to protein concentration and fluctuations in nutrients, temperature, and salt. Observations of context dependent multivalency in live cells suggest that both SpmX and PopZ form condensates *in vivo*.

Next, we investigated the relative topology and mixing behavior of SpmX and PopZ, as they could influence DivJ localization. Super-resolution microscopy of samples containing purified PopZ and SpmX at a physiological protein ratio exhibited SpmX clusters within 96% of PopZ condensates (Fig. 2F, top, materials and methods). However, PopZ condensates incubated with SpmXΔIDR, exhibited a uniformly distributed SpmXΔIDR signal (Fig. 2F, bottom). Cells over-expressing PopZ and harboring a dL5 fusion of SpmX (*19*) as the sole copy expressed at endogenous levels, exhibited multiple SpmX polar clusters in 89% of WT cells (Fig. 2G, top) and in 15% of SpmXΔIDR cells (Fig. 2G, bottom). These data suggest that the SpmX-IDR is necessary for driving SpmX self-association within PopZ condensates. *In situ* imaging of a fluorescent fusion of SpmX in *Caulobacter* cells at cryogenic temperatures (materials and methods) followed by overlaying the signal on the annotated ribosome excluded region (*20*), a proxy for the location of PopZ (*5*), revealed SpmX localizations clustered on one side of the PopZ microdomain (Fig. 2H). While the *in situ* images are from a small number of cells owing to a limited throughput of correlative imaging, the superior precision (∼5-10 nm) in SpmX localizations supports the notion that SpmX and PopZ coexist as two interacting, yet demixed condensates at the *Caulobacter* stalked pole.

Cumulatively, these data show that both SpmX and its localization determinant, PopZ can form phase separated condensates via multivalent interactions. Phase separation of PopZ and SpmX is a plausible mechanism enabling polar sequestration of client proteins (*14*), including DivJ thereby enhancing DivJ’s density dependent kinase activity. However, SpmX phase separation alone cannot explain the glucose dependent modulation of DivJ activity by SpmX-IDR (Fig. 1E). We hypothesized that SpmX multivalency could be sensitive to a glucose dependent metabolite. Since ATP is a key product of glucose metabolism, and has been shown to disaggregate eukaryotic condensates at high concentrations *in vitro* (*3*), we next focused on the role of ATP on SpmX and PopZ condensate formation.

### Solutes tune condensate multivalency in a protein-specific manner

Biomolecular condensates that contain disordered proteins can regulate their multivalency by interacting with ligands in their environment (*21*). Therefore, studying the phase behavior of condensates in the context of biologically relevant ligands can elucidate the nature of multivalent interactions. While 1,6-Hexanediol (1,6-HD) has been widely used to differentiate between material properties of condensates (*22*), neither is its mechanism of action is understood, nor is it produced in cells. More biologically relevant ligands such as ATP and Lipoic acid (LA) have been reported to dissolve eukaryotic protein condensates (*3, 23*). Accordingly, we sought to characterize SpmX and PopZ multivalent interactions in the presence of ATP, LA, and 1,6-HD.

Microscopic observation of sparsely labeled condensates incubated with increasing ATP concentrations showed that both PopZ and SpmX condensates dissolved in the presence of 1 mM, and 2 mM ATP, respectively (Fig. 3A, materials and methods). Notably, the dissolution concentrations of ATP for SpmX and PopZ were 5-10 fold lower than those observed for eukaryotic condensates (> 5 mM) (*3*) but spanned the physiological range of intracellular ATP for bacteria (*24*). In the presence of 1 mM ATP, SpmX formed metastable, hollow condensates that rapidly fused and dissolved (Video S5). Sparsely labeled SpmX and PopZ condensates dissolved upon LA treatment, with PopZ dissolution achieved at lower LA concentrations compared to SpmX (Fig. 3B). While LA is a cofactor for oxidative metabolism in oligotrophic bacteria (*25*), it is bound to proteins and not available freely in cells. Thus, cellular concentrations of LA may not be sufficient for condensate dissolution *in vivo*. Next, we tested the effect of the aliphatic alcohol 1,6-HD, on SpmX and PopZ condensates and found that PopZ condensation was promoted in a 1,6-HD concentration dependent manner (Fig. 3C), while SpmX condensation was inhibited under the same conditions. PopZ multivalency is likely to be promoted by 1,6-HD through the hydrogen bond acceptor residues (Proline, Aspartic acid, and Glutamic acid) present in its IDR (AA 1-102) and a continuous stretch of hydrophobic residues (AA 106-127) (Fig. S2B, domain diagram).

**Fig. 3.**
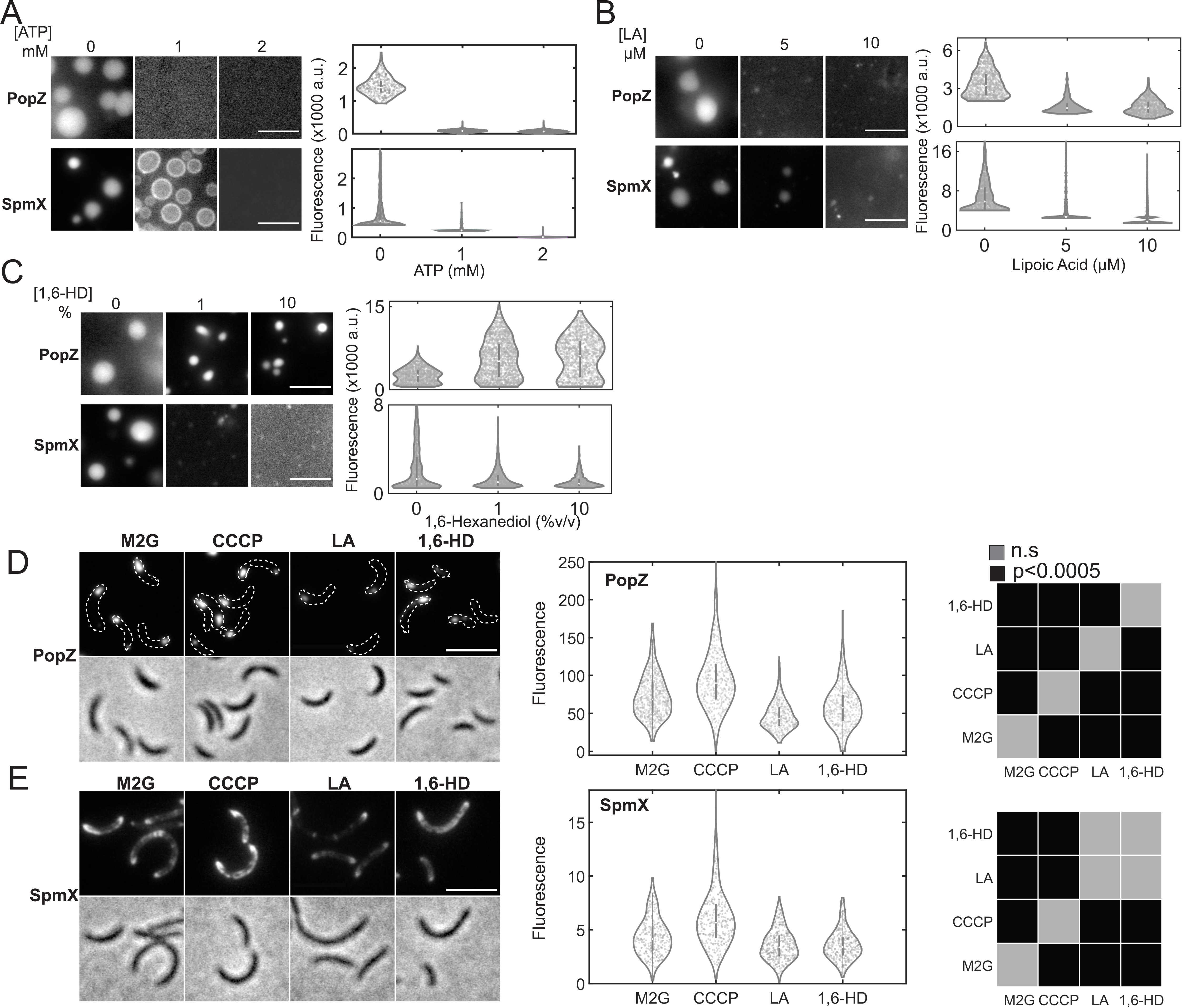
Solutes tune condensate multivalency in a protein-specific manner. (All scale bars unless noted, are 5 µm) **(A)** Fluorescence micrographs showing 5 µM PopZ (top, 1% Atto488 labeled) or 5 µM SpmX (bottom, ΔTM, 5% Cy3 labeled) incubated with indicated ATP concentrations for 1 hour in a physiological buffer (50 mM HEPES-KOH, pH 7.4, 0.1 M KCl, 5 mM MgCl_2_). Violin plots on the right show the distribution of the difference in fluorescence signal within the condensate and that outside the condensate, divided by the average fluorescence outside the condensate from a field of view. ∼600-1100 condensates were analyzed for each non-dissolving condition across three biological replicates. ATP can dissolve PopZ and SpmX condensates *in vitro*. **(B)** Fluorescence micrographs showing 5 µM PopZ (top, 1% Atto488 labeled) or 5 µM SpmX (bottom, ΔTM, 5% Cy3 labeled) incubated with indicated LA concentrations for 30 mins in a physiological buffer (50 mM HEPES-KOH, pH 7.4, 0.1 M KCl). For the case without LA, samples were incubated with an equivalent volume of the solvent for LA (DMSO). Violin plots on the right show the distribution of the difference in fluorescence signal within the condensate and that outside the condensate, divided by the average fluorescence outside the condensate from a field of view. ∼700-1100 condensates were analyzed for each non dissolving condition across three biological replicates. LA can dissolve PopZ and SpmX condensates *in vitro*. **(C)** Fluorescence micrographs showing 5 µM PopZ (top, 1% Atto488 labeled) or 5 µM SpmX (bottom, 5% Cy3 labeled) incubated with indicated 1,6-HD concentrations for 30 mins in a physiological buffer (50 mM HEPES-KOH, pH 7.4, 0.1 M KCl). Violin plots on the right show the distribution of the difference in fluorescence signal within the condensate and that outside the condensate, divided by the average fluorescence outside the condensate from a field of view. ∼600-1500 condensates were analyzed for each condition across three biological replicates. 1,6-HD promotes PopZ condensation while dissolving SpmX condensates *in vitro*. **(D)** Fluorescence and phase contrast micrographs of *Caulobacter* cells over-expressing eYFP-labeled PopZ, untreated (M2G) or treated with 100 µM CCCP (10 mins), 5 µM LA or 5% (v/v) 1,6-HD (30 mins, each). Middle panel shows violin plots with distribution of the ratio of localized to diffuse fluorescence within the cell (N∼900 cells per condition). Right panel shows pairwise statistical significance (p values) calculated for each distribution using a two sample t-test. *In vivo*, PopZ multivalency was promoted under ATP depletion and inhibited by LA treatment. 1,6-HD treated cells showed minimal difference from untreated cells. **(E)** Fluorescence and phase contrast micrographs of *Caulobacter* cells harboring a *popZ* deletion and over-expressing eYFP-labeled SpmX, untreated (M2G) or treated with 100 µM CCCP (10 mins), 5 µM LA or 5% (v/v) 1,6-HD (30 mins, each). Middle panel shows the distribution of the ratio of localized to diffuse fluorescence within the cell (N∼700 cells per condition). Right panel shows pairwise statistical significance (p values) calculated for each distribution using a two sample t-test. SpmX multivalency was promoted under ATP depletion and inhibited by LA and 1,6-HD treatment *in vivo*.

The intracellular environment may affect fluctuations in ligand concentrations, necessitating investigation of the observed *in vitro* stimulus responses in living cells. Accordingly, we tested the effect of rapid ATP depletion by the ionophore Carbonyl cyanide m-chlorophenyl hydrazone (CCCP) (Fig. S3A), or exposure to LA and 1,6-HD on PopZ and SpmX assemblies in live *Caulobacter* cells. Cells over-expressing eYFP-PopZ (Fig. 3D) or SpmX-eYFP (Fig. 3E) were treated with 100 µM CCCP (10 mins), 5 µM LA, or 5% 1,6-HD (30 mins, each) (materials and methods). Using the ratio of fluorescence from protein clusters to cellular background as a proxy for degree of condensation, we observed that depletion of ATP by CCCP addition promoted condensation for both PopZ and SpmX *in vivo*. While LA inhibited condensation for both proteins (Fig. 3D-3E), 1,6-HD treatment depleted only SpmX condensation. These data demonstrate solute dependent multivalency in SpmX and PopZ clusters, further supporting that clusters observed in live cells are phase separated condensates.

As a negative control, over-expression of DivJ-eYFP from a high copy plasmid in a *spmX* deletion strain led to the formation of DivJ clusters at polar and non-polar locations (Fig. S3B). However, unlike SpmX or PopZ condensates, DivJ-eYFP clusters exhibited a mild reduction in signal under all solute treatments. As another control we measured the effect of the ligands on the self-assembly of the structured intermediate filament protein Crescentin, which forms an extended filament on the dorsal side of *Caulobacter* cells (*26*). Like DivJ, CreS-eYFP fibers exhibited no change in fluorescence upon CCCP treatment, but they exhibited a slight increase in fluorescence upon LA or 1,6-HD treatment (Fig. S3C). Finally, as a positive control, we treated cells expressing RnaseE-eYFP, a key component of the *Caulobacter* RNA degradosome organelle (BR-bodies) (*27*) with CCCP, LA and 1,6-HD. RnaseE-eYFP condensation was promoted by ATP depletion, and 1,6-HD treatment, while being inhibited under LA treatment, compared to untreated cells (Fig. S3D). To assess the effect of chemical perturbations on cell health, we used a cell impermeant fluorescent dye as a proxy for membrane leakage (materials and methods). While LA and CCCP treated samples showed a similar number of cells with compromised membranes compared to untreated samples (∼1-2%), HD treated samples exhibited ∼8% cells with membrane leakage (Fig. S3E). The effects of 1,6-HD on membrane integrity shown here, and enzyme activity reported previously (*28*) underscore the pleiotropic consequences of the compound *in vivo*.

Cumulatively, the data show that phase separation within three different *Caulobacter* condensates is promoted at low ATP levels and inhibited by LA, whereas structured protein assemblies (CreS) and protein-rich aggregates (DivJ over-expression) do not exhibit such a behavior. While 1,6-HD inhibits SpmX condensation, it promotes PopZ condensation likely by promoting multivalent interactions via hydrophobic residues and hydrogen bond acceptor moieties. Critically, the data challenge the widely held dogma that molecules such as 1,6-HD generally inhibit phase separation while supporting the notion of protein and ligand specific effects on multivalency, phase separation and cell membrane integrity. Having provided evidence that ATP modulates SpmX multivalency, we asked whether ATP’s role as a solute is relevant for DivJ kinase regulation via the SpmX-IDR.

### Interplay between ATP and SpmX multivalency modulates DivJ activity

Given that DivJ exhibited density dependent kinase activity (Fig. S1E), we asked if under low intracellular ATP concentrations, the promotion of SpmX condensation leads to enhanced DivJ polar sequestration and kinase activity. Accordingly, we measured if DivJ exhibited ATP dependent polar localization in strains bearing WT SpmX. Cells over-expressing SpmX-eYFP and DivJ-mCherry in a *popZ* deletion strain exhibited enhanced DivJ polar signal under CCCP triggered ATP depletion (Fig. S4A), a phenotype that was not observed in the absence of SpmX (Fig. S3B). Next, we tested whether ATP-dependent DivJ sequestration via SpmX-IDR led to a metabolic control of cell division in stalked cells. We used aberrant cell division as a proxy for DivJ activity as before (Fig. 1E), while growing cells in minimal media with varying glucose concentrations. Cells grown under a five-fold lower glucose concentration for four hours exhibited ∼ 18% decrease in ATP levels (Fig. S4B), measured using a commercial Luciferase based assay (materials and methods).

In the WT strain, the number of aberrantly dividing cells increased with higher intracellular ATP concentrations (Fig. 4A, materials and methods). A DivJ kinase mutant strain exhibited ∼ten-fold higher number of filamentous cells compared to WT cells. Interestingly, the number of observed filamentous cells in DivJ kinase mutant was proportional to ATP levels, as observed for the WT strains. In stark contrast to WT or DivJ kinase mutant cells, SpmXΔIDR cells grown under high glucose phenocopied WT cells, and under low glucose phenocopied the DivJ kinase mutant (Fig. 4A). This phenotype could be rescued by mild expression of a WT copy of *spmX* under the control of a chromosomal Xylose promoter in SpmXΔIDR cells (Fig. 4A and S1I). The notion of ATP-dependent DivJ modulation was also supported by cell growth assays, where the rate of increase in cellular biomass was three times slower for SpmXΔIDR cells compared to WT cells under low glucose concentrations (Fig. 4B).

**Fig. 4.**
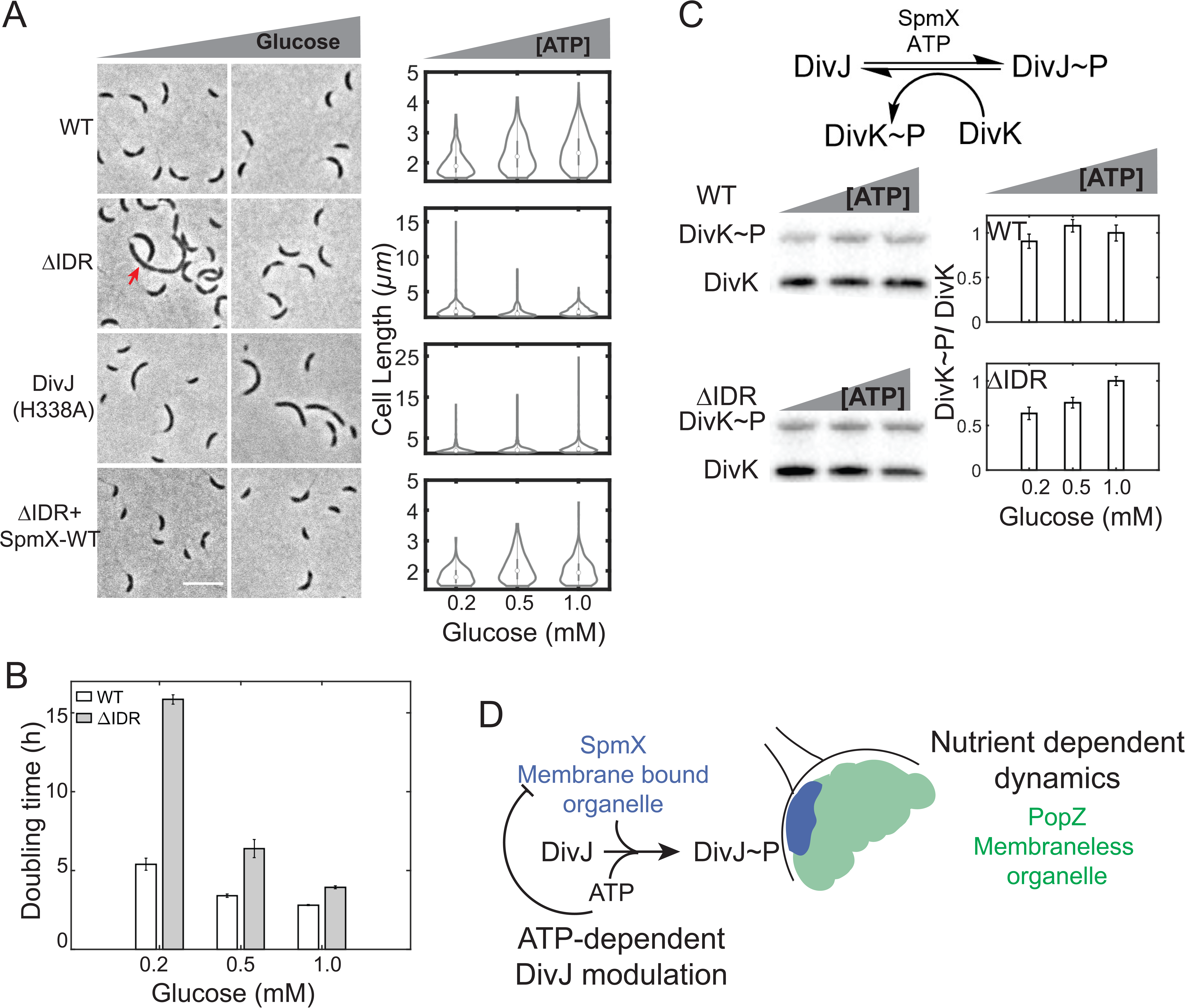
Interplay between ATP and SpmX multivalency modulates DivJ activity. **(A)** Representative phase contrast images (from top to bottom) of WT cells, SpmXΔIDR cells, DivJ(H338A) cells, and SpmXΔIDR cells with mild expression of WT SpmX, grown in M2 minimal medium with varying amounts of glucose (0.2 mM (left) to 1 mM (right)). Red arrow in the ΔIDR strain imaged under low glucose condition denotes an aberrantly dividing cell. Violin plots on the right show the distribution of lengths for the respective strains as a function of increasing glucose and intracellular ATP. The relative decrease in ATP concentrations across the range tested is ∼18%. Note the difference in the ordinate for WT (1-5 µm), SpmXΔIDR (1-15 µm) and DivJ kinase mutant (1-25 µm). SpmXΔIDR exhibits an ATP-dependent phenotype, phenocopying WT cells under high ATP, and the DivJ kinase mutant under low ATP concentrations. Mild expression of WT SpmX in cells harboring SpmXΔIDR rescues the aberrant division phenotype (Scale bar 5 µm). **(B)** Cell growth assayed through doubling time as a function of glucose in the growth medium for WT and SpmXΔIDR cells. Glucose concentration in M2 media is denoted on the abscissa. Doubling times were calculated by averaging at least three time lags during which the optical density measured at 600 nm doubled in the exponential growth phase. Each measurement was performed in triplicate. Error bars represent standard error of the mean from three biological replicates. SpmXΔIDR cells exhibited a slower growth rate compared to WT cells as glucose was depleted. **(C)** (Top) Schematic showing the determinants of DivJ kinase reaction. SpmX-IDR and ATP promote DivJ phosphorylation. DivK is a response regulator that dephosphorylates DivJ. (Bottom, Left) PhosTag SDS-PAGE immunoblots show the phosphorylation of DivK in extracts from stalked and pre-divisional WT cells (top) or SpmXΔIDR cells (bottom) as a function of intracellular ATP levels. DivK was fused to a Flag peptide and an anti-Flag antibody was used to detect both phosphorylated and unphosphorylated forms of DivK. (Bottom, right) Bar plots show the ratio of phosphorylated and unphosphorylated DivK measured from three replicates of the PhosTag assay. DivK (and by extension, DivJ) phosphorylation is robust to ATP fluctuations in WT cells, while being proportional to ATP levels in SpmXΔIDR cells. **(D)** Model showing the relative topologies of the interacting condensates formed by SpmX and PopZ. Based on our observations, both PopZ and SpmX exhibit nutrient dependent dynamics, and SpmX renders an ATP dependent feedback on DivJ regulation, resulting in robust modulation of DivJ even under low ATP concentrations.

To test whether aberrant cell divisions in SpmXΔIDR cells were indeed a result of reduced DivJ kinase activity, we sought to measure the levels of phosphorylated DivJ in lysates from cells using PhosTag gel electrophoresis as a function of decreasing glucose in the growth media (materials and methods). Owing to the low copy numbers of DivJ (∼12 molecules/ pole) and the labile nature of phospho-histidine, we were unable to robustly measure DivJ∼P levels using PhosTag assay. Therefore, we relied on DivJ’s response regulator DivK as a proxy for DivJ∼P levels in stalked and pre-divisional cells. DivK dephosphorylates DivJ (Fig. S4C) and forms a relatively stable phospho-aspartate bond allowing a robust measurement via PhosTag. WT and SpmXΔIDR cells expressing an endogenous Flag-tagged copy of DivK were grown under decreasing glucose concentrations. Lysates from these cells were subjected to PhosTag gel electrophoresis, where phosphorylation dependent protein migration enables separation between phosphorylated and unphosphorylated forms of the same protein. PhosTag gels were immunoblotted and both forms of Flag-tagged DivK were detected using an antibody against Flag-tag. While the DivK∼P/DivK ratio correlated with the decrease in glucose (ATP) for SpmXΔIDR cells, such an ATP-dependence was not observed in WT cells (Fig. 4C). These data show that ATP and SpmX-IDR dependent regulation of cell length is due to changes in DivJ phosphorylation levels and underscore the importance of SpmX-IDR in promoting DivJ activity under ATP depletion. Cumulatively, the data establish SpmX as a membrane associated condensate that compartmentalizes DivJ and promotes its kinase activity. The sensitivity of SpmX condensation to physiological intracellular ATP levels acts as a feedback to enhance DivJ cooperativity, particularly when its substrate ATP is scarce, thereby enabling a robust modulation of DivJ activity.

## Discussion

Cellular processes critical for survival and adaptation must be robust across a broad range of nutrient conditions. Unlike growth in lab media, bacteria in their natural habitats experience large fluctuations in nutrient concentrations (*29*) and must adapt their biochemistry for survival particularly under nutrient depletion. Yet, mechanisms that connect metabolic state of the cell to its biochemistry are poorly understood. Here, we investigated the mechanisms that impart robustness to the DivJ kinase under ATP limitation in the aquatic bacterium, *Caulobacter crescentus*. Our data show that DivJ polar localization and kinase activity are promoted by the disordered domain of SpmX. SpmX, and its localization determinant, PopZ form interacting, but demixed condensates through multivalent interactions. Chemical probing of SpmX and PopZ condensates revealed solute and protein specific multivalency. While it is not possible to ascertain the specific phase behavior of SpmX and PopZ in living cell due to size constraints, the observation of context dependent multivalency supports the idea that these proteins exist as biomolecular condensates *in vivo*. Critically, the data show that ATP depletion promoted multivalency in SpmX and PopZ condensates *in vivo*. The ATP-dependent condensation of SpmX conferred a feedback on DivJ sequestration, promoting DivJ kinase activity under low ATP concentrations. Our observations explain the nutrient dependent dynamics of PopZ and underscore the relevance of SpmX-IDR in DivJ regulation under low glucose conditions (Fig. 4D).

While certain eukaryotic cells and symbiotic bacteria often thrive in relatively stable environments, free-living bacteria are constantly exposed to chaotic cycles of famine and feast. Many enzymes critical for cell proliferation are present at low copy numbers, which poses an additional challenge for cells surviving under substrate depletion. Condensates formed by multivalent interactions among disordered proteins enable selective sequestration of low copy number enzymes (*14*) and buffer against the noise associated with stochasticity in enzyme activity (*30*). Our results suggest that compartmentalization of a low copy number kinase within an ATP-sensitive condensate is an efficient mechanism to further reduce the biochemical noise under nutrient depletion. Notably, such a mechanism enables an ATP-dependent degree of cooperativity of an enzyme, as opposed to a fixed degree of cooperativity (such as the Hill coefficient) and provides a boost to enzyme activity under substrate limitations. Further, due to faster diffusion of ATP compared to gene expression time scales, condensates sensitive to intracellular-ATP concentration range may adapt swiftly and robustly to changing metabolic conditions.

In eukaryotic cells, high levels of intracellular ATP (5-10 mM) have been attributed to the maintenance of a soluble proteome. Why are bacterial condensates soluble within a much lower ATP range (0.5-1 mM) compared to eukaryotic condensates? It has been observed that the intrinsic disorder content of a proteome is correlated with cellular complexity (*31*). Thus, we posit that the emergent ATP-dependent multivalency in disordered proteins has been preserved through evolution due to a concomitant increase in cellular complexity and energy consumption. This idea is also supported by recent observations of ATP dependent protein condensation (*32*) and aggregation (*33*) in bacteria. ATP has been shown to affect the diffusion of mesoscopic macromolecular assemblies in bacteria (*34, 35*). We show that multivalency modulation by ATP affects molecular diffusion even within mesoscopic condensates and may be a critical mechanism for optimal functioning of compartments formed by disordered proteins. Thus, we propose that the diverse repertoire of condensates can serve as control knobs to tune multivalency and reactivity in response to the metabolic state of the cell.

## Notes

## Acknowledgments

The authors thank Dr. Rohit Pappu, Dr. Jared Schrader, Dr. Sean Crosson and Dr. Sunish Radhakrishnan for helpful discussions on experimental aspects of the manuscript. The authors also thank Drs. Harley McAdams, Richard Losick, Kotaro Fujii, Dante Ricci, Edward de Koning and Bo Gu for a careful reading of the manuscript. The authors thank all members of the Shapiro and Moerner labs for their constructive feedback throughout this work.

## Author contributions

This study was conceived by S.S. Experiments were designed by S.S., with inputs from W.E.M. and L.S. Strains for this study were engineered by S.S. and T.N.C. Single molecule imaging and data analyses were performed by C.B. and S.S., with inputs from W.E.M. Confocal imaging was performed by H.N.C. and S.S. Electron microscopy was performed by P.D.D. and S.S. The manuscript was written by S.S. and L.S., with inputs from W.E.M.

## Data and materials availability

All data is available in the main text or the supplementary materials. All data, code, and cell strains used in this work are available upon request.

## Supplementary Materials

Supplementary Figures - Figs. S1-S4

Captions for Movies

Supplementary text

Materials and Methods

Movies S1 through S5

## Supplementary text

### Assessment of DivJ diffusion using mean squared displacement

To further validate CDF assessments of DivJ trajectories, mean-squared-displacement (MSD) analyses (*14*) were performed. MSD analyses also indicated that DivJ diffusion was on average ten-times faster in the polar microdomain in the absence of SpmX-IDR (Fig. S1C). As a negative control, we engineered a Halo enzyme fused to the trans-membrane helical region from an *E. coli* sensor protein ArcB, such that the resulting fusion does not interact with SpmX or any other proteins in *Caulobacter*. The membrane-localized Halo enzyme displayed the same diffusivity in the pole in the presence or absence of SpmX (Fig. S1D), indicating that the differences in diffusion observed for DivJ are specific to its interaction with SpmX. Cumulatively, single molecule diffusion analyses indicate that DivJ is localized to the stalk-bearing pole through an interaction that is facilitated by the SpmX-IDR, resulting in a higher polar dwell time and sequestration of DivJ.

### eYFP fusion of SpmX-IDR

SpmX, on account of containing structured and disordered domains, exhibits oligomerization and multivalency that is dependent on both its domains (*7*). Accordingly, we observed that SpmX-IDR (AA156-355) was unstable in E. coli and as a result could not be purified to concentrations above 5 µM. Likely, this was a result of the SpmX-IDR getting proteolyzed in E. coli due to its exposed disordered termini. Several strategies were used to “cap” the N-terminus of the IDR, including His-Tag, Flag-Tag and eYFP-fusion. While the His-tagged and Flag-tagged proteins exhibited meagre improvements in yield, the eYFP fused IDR could be purified at high concentrations (∼20-25 µM). Despite the improved yield eYFP-IDR condensates were unstable under imaging conditions, and they collapsed and degraded upon touching the glass surface. As a result, they were imaged while diffusing above the glass surface. This imposed a challenge in two-color imaging of eYFP-IDR and DivJ (sparsely labeled with Cy3), which was overcome by 3D imaging of eYFP-IDR condensates in the presence of DivJ (Movies S1-2). Enrichment of DivJ signal was observed in all eYFP-IDR condensates, indicative of a direct interaction between SpmX-IDR and DivJ *in vitro*. This observation could also explain the SpmX-IDR dependent dwell time exhibited by DivJ in the polar microdomain, measured via single protein tracking.

### PopZ gelation states

Upon imaging PopZ that was reconstituted directly from frozen protein stocks, we observed non-spherical condensates reminiscent of a “beads on a string” morphology. These morphologies likely represented thermally trapped states. The beads on a string morphology of PopZ condensates could be relieved using heat treatment. While both forms of PopZ showed arrested dynamics *in vitro*, PopZ within the annealed condensates exhibited slower internal rearrangements compared to untreated condensates suggesting that annealing promoted multivalent self-interactions (Fig. S2I). Accordingly, untreated PopZ condensates ripened while annealed PopZ condensates fused albeit over similar time scales (Fig. S2J). These data suggest that PopZ exhibits different states exhibiting slow internal dynamics and separated by a thermal barrier *in vitro*.

## Figures – Supplementary text

**Fig. S1.**
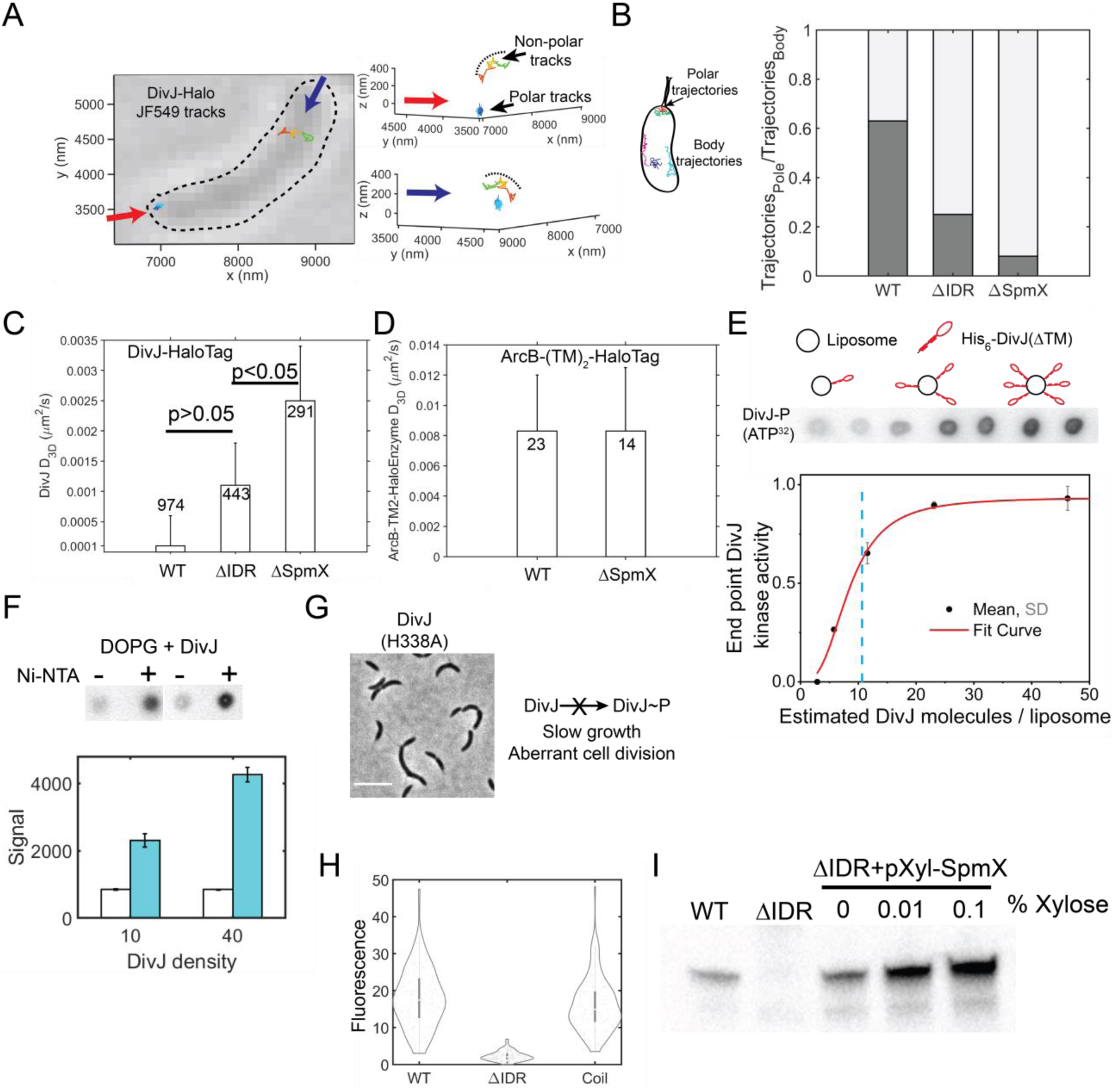
Effect of SpmX-IDR on DivJ localization, diffusion, and activity. Throughout this panel, ΔIDR denotes SpmXΔIDR. **(A)** (left) Representative bright-field image of a live *Caulobacter* cell, overlaid with 2D projections of 3D DivJ-HaloTag trajectories measured using single-particle tracking. Each trajectory is plotted using a different color. Arrows denote perspectives for the 3D representation shown on the right. (right) Representation of the trajectories from the left panel in 3D space with appropriate perspective and all three Cartesian axes. Green and red trajectories in the body are membrane-associated as can be seen from their curvature, highlighted using a black dashed line. 3D tracking reveals polar as well as non-polar membrane associated trajectories. **(B)** Fraction of single molecule DivJ trajectories observed in the pole vs the body for respective strains. Total tracks analyzed are 1558 (WT), 1764 (ΔIDR) and 3572 (ΔSpmX). SpmX-IDR enables DivJ sequestration at the pole. **(C)** Three-dimensional diffusion coefficients obtained from Mean Squared Displacement (MSD) analyses for polar DivJ trajectories in various SpmX backgrounds. The number of trajectories analyzed for each sample are noted with the bars. P values from a pairwise two-tailed t-test are denoted in the plot. **(D)** 3D diffusion coefficients obtained from MSD analyses of control trans-membrane HaloTag trajectories in WT and SpmX deletion backgrounds. The HaloTag was attached to the membrane using two trans-membrane helices from the *E. coli* protein ArcB. Differences were not detected in the polar diffusion coefficients of the HaloTag in WT or SpmX deletion backgrounds. Errors in D in both panels (C) and (D) are standard deviations estimated from bootstrapping (50% sampling with replacement performed 500 times). **(E)** DivJ auto-kinase activity assayed on liposomes. Liposomes contained 90% (mol %) di-oleoyl-phosphatidyl glycerol (DOPG) and 10% (mol %) Nickel-chelated lipids (DGS-NTA) to bind His_6_-DivJ(ΔTM). A fixed concentration of 5 μM His_6_-DivJ(ΔTM) was pre-incubated with increasing amounts of liposomes to obtain increasing surface density of His_6_-DivJ(ΔTM) molecules per liposome. Kinase reaction was performed in a buffer (25 mM HEPES-KOH, pH 7.4, 50 mM KCl, 25 mM NaCl, 5 mM MgCl_2_) by adding 0.1 µCi ATP(γ−^32^P) for 5 min, quenching and blotting on a nitrocellulose membrane followed by phosphor imaging. The measured activity at each His_6_-DivJ(ΔTM) surface density state was normalized to the highest density condition. Representative phosphor imaging data from spot assays and a schematic depicting increasing DivJ density (red) on liposomes (black circles) are shown above the graph. Intensity from phosphor imaging spots was quantified and the average data from triplicate spot assays were fit to a Hill-Langmuir curve as described in the methods section. Dashed blue line represents the average physiological polar DivJ concentration of 11 molecules measured from a single molecule counting assay for cells grown in M2G media. Error bars represent the standard deviation from 3 measurements. DivJ exhibited a density dependent kinase activity *in vitro*. **(F)** Liposome phosphorylation assay with (cyan bars) or without (white bars) Ni-NTA lipids at two different estimated densities of 10 and 40 DivJ molecules/ liposome. Error bars are standard deviations from 3 measurements. DivJ signal from liposomes without Ni-NTA lipids is insensitive to the protein density. **(G)** Representative phase contrast image of DivJ (H338A) cells in which DivJ phosphorylation is absent, leading to aberrant cell division (scale bar 5µm). **(H)** Distribution of the ratio of mean polar and non-polar DivJ signal in WT cells (N=235), ΔIDR cells (N=258) and cells with SpmXΔIDR fused to a CoilZ peptide (N = 204) that can bind to a cognate CoilY peptide fused to DivJ-eYFP. Coil tag recruitment of DivJ gives a polar signal comparable to WT cells independent of the SpmX-IDR. **(I)** Western blot showing the levels of WT SpmX in WT, SpmXΔIDR, and SpmXΔIDR cells expressing WT SpmX on a chromosomal Xylose promoter (rescue cell line). For the rescue cell line, WT SpmX was produced by adding the indicated amount of Xylose. Western blot was performed using an antibody raised against the IDR of SpmX (*6*). Even without Xylose addition, the mild expression of SpmX was comparable to SpmX levels in WT cells.

**Fig. S2.**
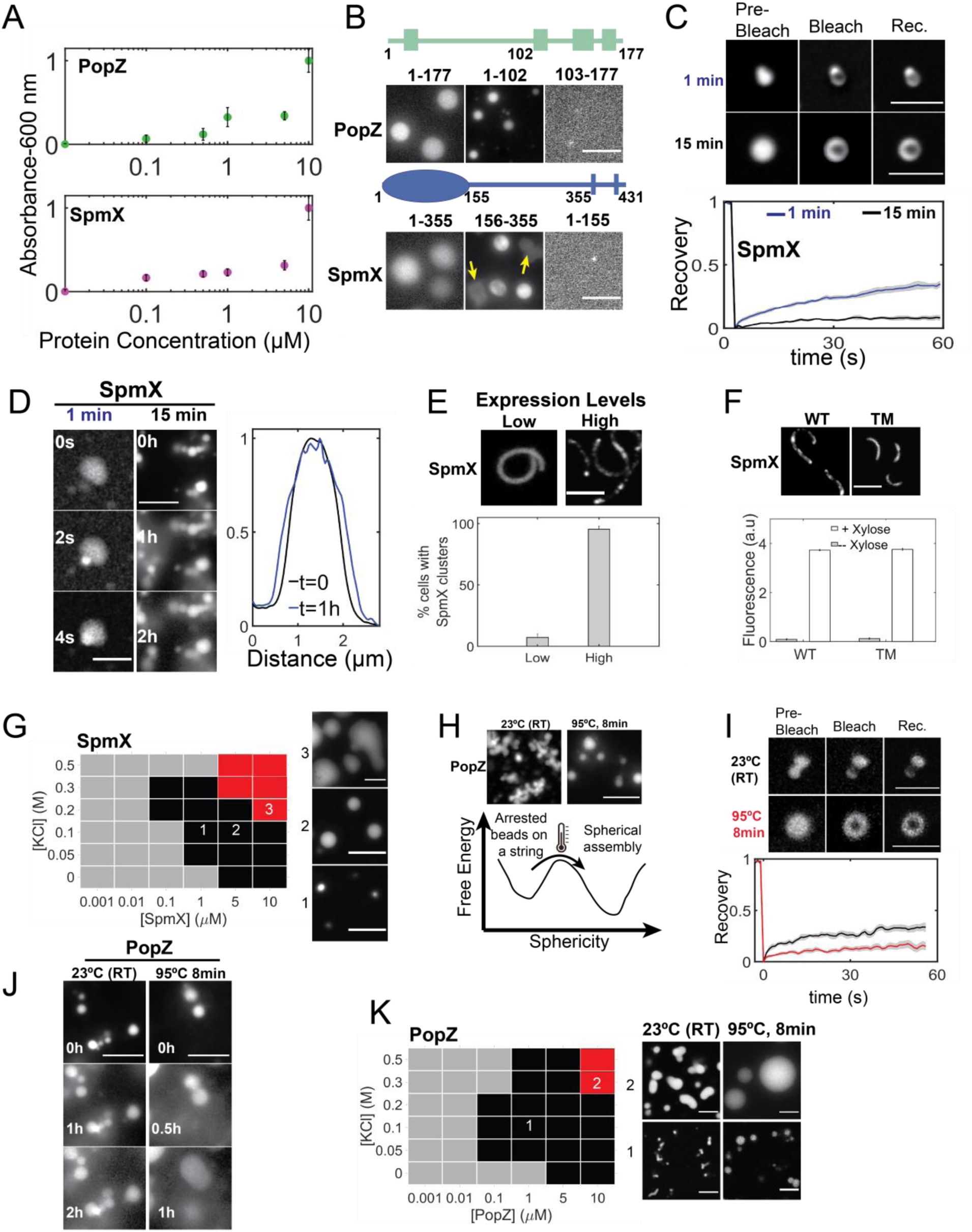
Context dependent properties of SpmX and PopZ condensates. (All scale bars in this figure are 5 µm.) **(A)** Solution turbidity measured as a function of increasing PopZ (top) or SpmX(ΔTM) (bottom) concentration in a physiological buffer (50 mM HEPES-KOH, pH 7.4, 0.1 M KCl) at room temperature. Background subtracted-absorbance values were normalized to the maximum absorbance observed in each case. Error bars represent standard deviation from 3 independent measurements. PopZ and SpmX exhibit a concentration dependent increase in turbidity. **(B)** (top) Representative fluorescent micrographs of PopZ domains: PopZ (full length), PopZ (AA 1-102) and PopZ (AA 103-177) (all proteins 1% Atto488 labeled). (bottom) Representative fluorescent micrographs of SpmX domains: SpmX (ΔTM, 5% Cy3 labeled), eYFP-tagged SpmX-IDR, and SpmXΔIDR (ΔTM, 5% Cy3 labeled). Yellow arrows in eYFP-tagged SpmX-IDR sample indicate condensates that degraded upon contacting the coverslip. All proteins are at 5 µM concentration. IDR containing regions of both SpmX and PopZ are necessary and sufficient for phase separation *in vitro*. **(C)** Internal rearrangement of SpmX condensates was assayed by FRAP at two different time points after fusing on glass. (top) A 250 nm spot was bleached in a SpmX condensate (5 µM protein, ΔTM, 5% Cy3 labeled), and the change in the fluorescence was analyzed as a function of time for condensates relaxed on glass within 1 min (blue) or after 15 mins (black). (bottom) Analyses of the fluorescence recovery of 1-min-old (black; N = 9), 15-min-old (green; N = 11) condensates are shown. Gray shadow depicts the standard error of the mean. SpmX condensates exhibit concentration dependent dynamics *in vitro*. **(D)** (Left) Condensates of SpmX fused to glass within 1 min (5 µM protein, ΔTM, 5% Cy3 labeled) displayed spontaneous fusion on the seconds time scale, and (Right) condensates older than 15 mins ripen on the hours’ time scale without fusion. Measurement of the diameter of a ripening SpmX condensate over 1 hour showed an increase in its diameter by ∼350 nm. Ripening is associated with slow condensate internal dynamics as a result of protein self-association. **(E)** Fluorescence micrographs of *Caulobacter* cells harboring a *popZ* deletion and over-expressing eYFP-labeled SpmX on a chromosomal Xylose promoter, induced using 0.03% Xylose (left) or 0.3% Xylose (right) for 1 hour. Bar graph below shows the % of cells exhibiting two or more SpmX clusters per cell (N ∼ 400 cells for each case). SpmX cluster formation is concentration dependent *in vivo*. **(F)** Representative fluorescence micrographs of cells expressing SpmX-eYFP in a *popZ* deletion (left) or SpmX(TM)-eYFP (Δ1-355) in a *spmX* deletion, on a chromosomal Xylose promoter, induced using 0.3% Xylose for 1 hour. While clusters were observed for WT SpmX (>95% cells), SpmX-TM (Δ1-355) always remained diffuse. Shown below are end point YFP fluorescence measured using a plate reader in cells expressing SpmX-eYFP or SpmX-TM-eYFP (Δ1-355) grown in the presence or absence of 0.3% Xylose for 1 hour. At comparable concentrations, SpmX-TM domain without the cytoplasmic domain was unable to form clusters *in vivo*. **(G)** Phase diagram of SpmX as a function of protein and KCl concentration. Black squares denote conditions under which >20 fluorescent spots were observed to be above the intensity threshold (<background> + 3σ_background_) from 80 µm square regions across six fields of view spanning two biological replicates. Red squares denote metastable condensates suggestive of the spinodal decomposition on the phase diagram. Representative fluorescent micrographs from conditions 1-3 are shown next to the plot. **(H)** Representative maximum intensity projections of 3D confocal micrographs of PopZ (5 µM, 1% Atto 488 labeled) reconstituted in a physiological buffer (50 mM HEPES-KOH, pH 7.4, 0.1 M KCl). PopZ condensates exhibited “beads on a string” morphology at 23 °C (left) but relaxed into spherical condensates upon heat treatment (right). This suggests the presence of a thermal barrier between conformational states of different sphericity, depicted in the schematic below. **(I)** Internal rearrangement of PopZ condensates (5 µM, 1% Atto488 labeled) *in vitro* was assayed by FRAP. (top) A 250 nm spot was bleached, and the change in the fluorescence was analyzed as a function of time for untreated condensates (black) and annealed condensates (red). (bottom) Analysis of fluorescence recovery of untreated condensates (black; N = 10), annealed condensates (red; N = 12) are shown. Gray shadow depicts the standard error of the mean. Annealed condensates exhibit slower internal dynamics compared to arrested condensates. **(J)** Representative fluorescence micrographs showing ripening behavior of arrested PopZ condensates (5 µM protein, 1% Atto488 labeled) without fusion on the hours’ time scale *in vitro*. Annealed PopZ condensates fused over sub-1 hour time scales showing the differences between the slow dynamic states of condensates *in vitro*. **(K)** Phase diagram of PopZ condensates (1% Atto488 labeled) as a function of protein and KCl concentration. Black squares denote conditions under which >20 fluorescent spots were observed to be above the intensity threshold (<background> + 3σ_background_) from 80 µm square regions across five fields of view spanning two biological replicates. Red squares denote metastable condensates suggestive of a spinodal decomposition on the phase diagram. Representative fluorescent micrographs from conditions 1-2 for the untreated and heat treated samples are shown next to the plot.

**Fig. S3.**
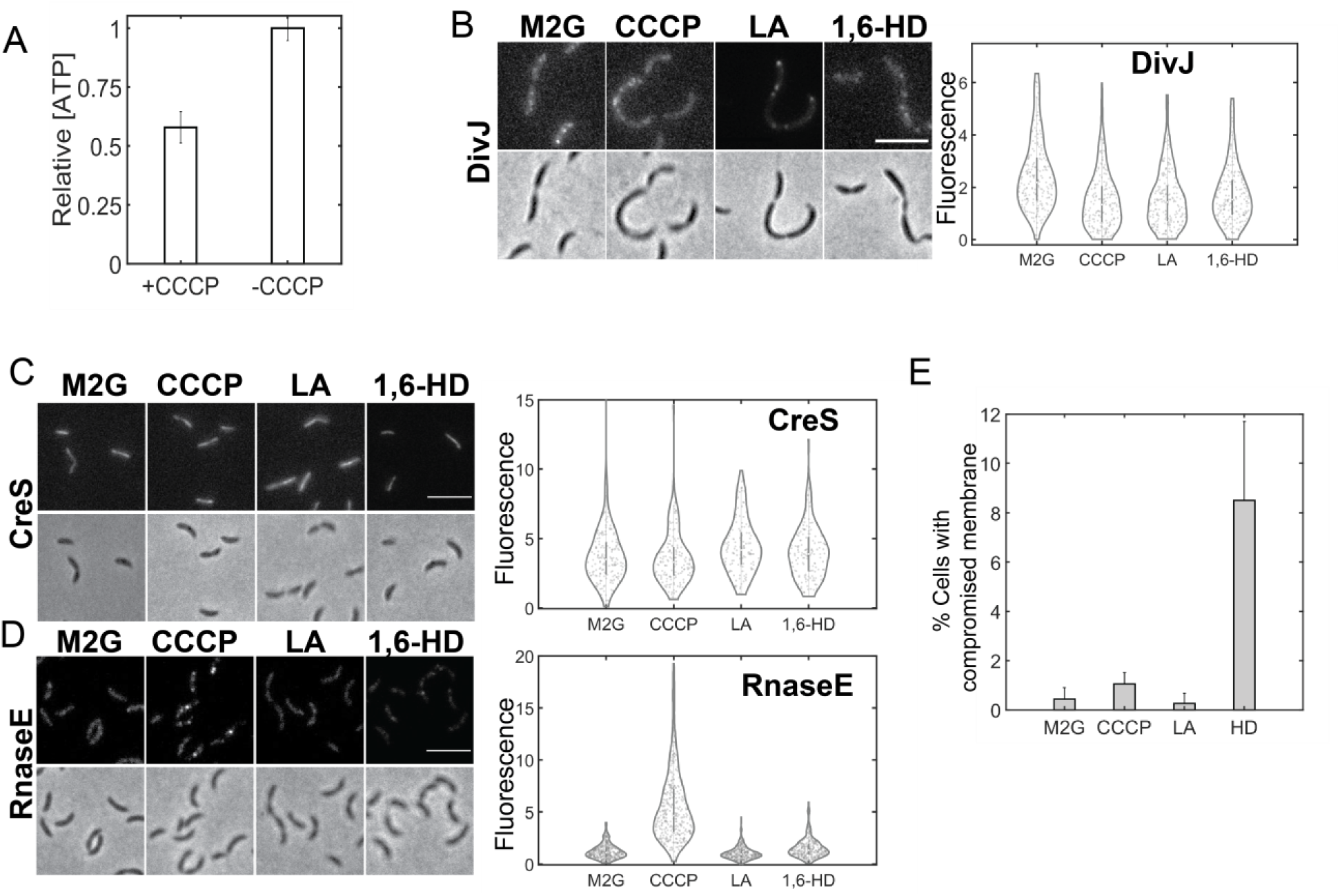
Effect of solutes on various protein assemblies and membrane integrity *in vivo*. (All scale bars in this figure are 5 µm.) **(A)** Relative intracellular ATP concentrations in *Caulobacter* cells treated with 100 µM CCCP dissolved in DMSO, or an equivalent volume of DMSO for 10 minutes, measured using a commercial Luciferase based assay. Background from the cell growth media was subtracted and luminescence units were normalized to the ATP levels in the DMSO (-CCCP) case. Error bars represent the standard deviation from 3 biological replicates, with each sample assayed in triplicate. CCCP addition leads to rapid ATP depletion in live *Caulobacter* cells. **(B)-(D)** Effect of ATP depletion by CCCP addition (100 µM, 10 min), 5 µM LA, or 5% (v/v) 1,6-HD treatment (30 mins, each) on various protein assemblies *in vivo*. **(B)** Representative fluorescence (top) and phase contrast (bottom) micrographs of *Caulobacter* cells with *spmX* deletion and over-expressing DivJ-eYFP on a high copy Xylose inducible plasmid exhibited fluorescent clusters. However, the fluorescence within DivJ-eYFP clusters was depleted under all treatments (CCCP, LA, 1,6-HD). ∼600 cells were analyzed for each condition. **(C)** Representative fluorescence and phase contrast micrographs of *Caulobacter* cells over-expressing Cres-eYFP exhibited fluorescent fibers. The relative fluorescence intensity in these fibers depleted under CCCP treatment, while a mild enhancement in fluorescence was observed under HD or LA treatment. ∼500 cells were analyzed for each condition. **(D)** *Caulobacter* cells expressing endogenously tagged RnaseE-eYFP exhibited clusters with enhanced fluorescence under CCCP treatment (10 mins) and depleted fluorescence under LA treatment but not under HD treatment (30 mins, each). ∼800 cells were analyzed for each condition. ATP and LA, but not HD, are able to dissolve condensates such as BR bodies while having different effects on structured protein assemblies such as CreS, or protein aggregates such as DivJ in the absence of SpmX *in vivo*. **(E)** Bar plot showing percentage of *Caulobacter* cells expressing eYFP-PopZ with membrane leakage observed under treatment with CCCP (N = 762 cells), LA (N=682 cells) and HD (N = 474 cells), compared with the untreated cells (M2G, DMSO, N = 719 cells). Cells were treated with the respective solutes followed by labeling with Live-or-Dye^TM^ (640/662) dye based on manufacturer’s protocols. Dye labeled cells were washed thrice using M2G and imaged on an epifluorescence microscope. Cells with compromised membranes exhibited at least five-fold higher fluorescence compared to healthy cells. This threshold criterion was applied to all the images to obtain the bar plots. Error bars are standard deviation across 12 fields of view from three biological replicates. 1,6-HD treated cells exhibited 4-5-fold higher membrane leakage than CCCP or LA treated cells.

**Fig. S4.**
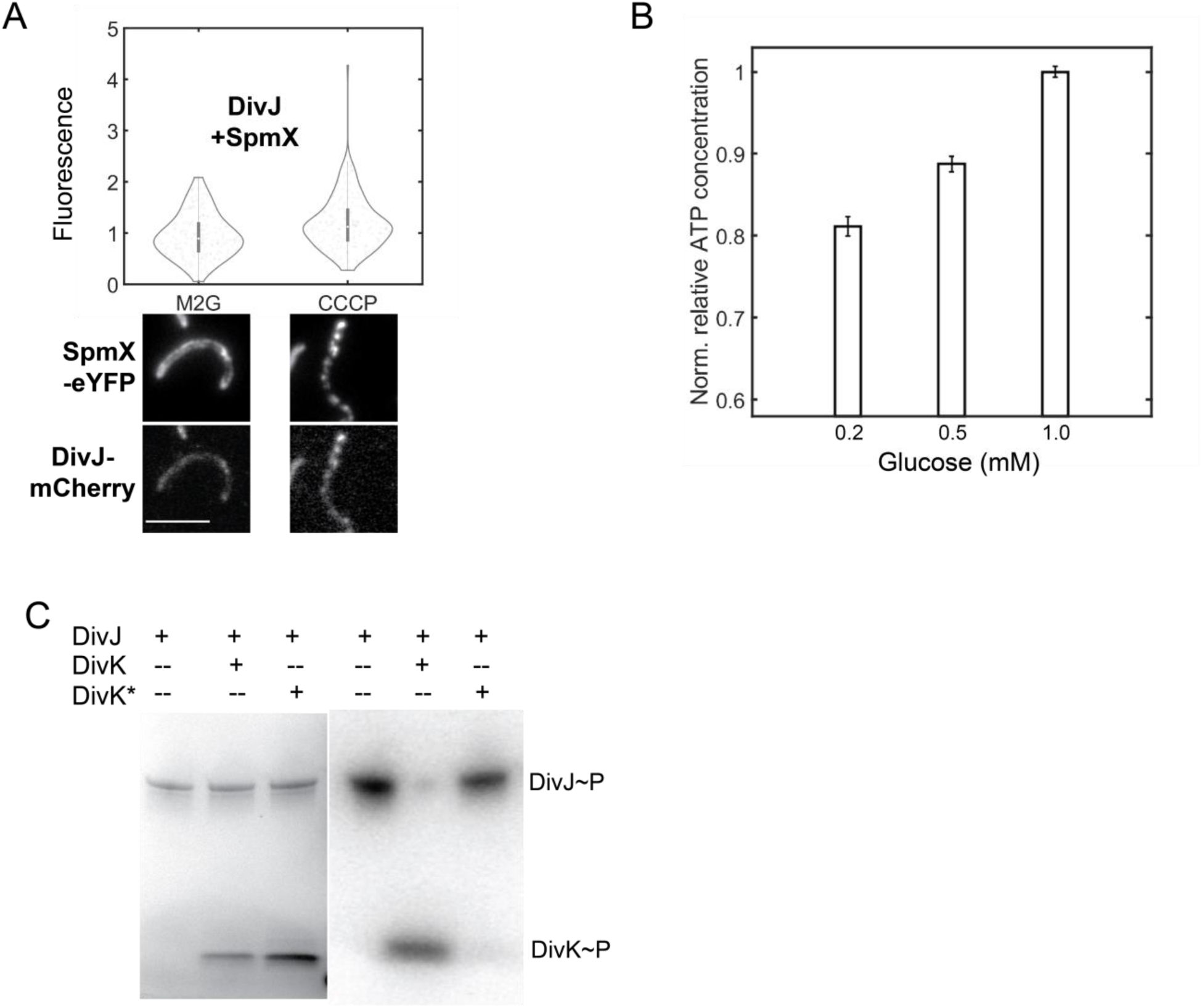
Effect of ATP on DivJ localization and phenotype. **(A)** Distribution of the ratio of polar localized to diffuse DivJ signal in *Caulobacter* cells co-expressing SpmX-eYFP and DivJ-mCherry in a strain harboring *popZ* deletion under the control of Xylose and Vanillate promoters, respectively. DivJ-mCherry was expressed using 50 mM Vanillate while SpmX-eYFP cluster formation was induced by the addition of 0.3% Xylose for 4 hours. Cells were treated with DMSO (M2G) or 100 µM CCCP dissolved in DMSO (10 mins) followed by imaging. Representative fluorescence micrographs of cells in both eYFP and mCherry channels are shown below (scale bar is 5 µm). ∼400 cells were analyzed for each condition. Polar localization of DivJ is enhanced under ATP depletion (scale bar 5µm). **(B)** Bar plot showing the relative intracellular ATP concentrations in *Caulobacter* cells grown in M2 minimal media as a function of glucose concentration from 0.2 mM to 1 mM. ATP concentrations were measured using a commercial Luciferase based luminescence assay. Error bars represent standard error of the mean from 3 biological replicates each assayed in triplicate. **(C)** *In vitro* phosphorylation measurement of the cytoplasmic domain of DivJ (5 µM) alone, in the presence of DivK (5 µM), or DivK* (DivK(D53N)) (5 µM) incubated with 1 µCi ATP(γ−^32^P) in a physiological buffer (50 mM HEPES-KOH, pH 7.4, 0.1 M KCl, 5 mM MgCl_2_) for 5 mins. Phosphorylation reactions were quenched by addition of SDS, and proteins were separated via SDS-PAGE. (left) Silver-stained SDS-PAGE gel with DivJ, DivJ and DivK or DivJ and DivK* (Right) Radiograph showing the phosphorylation levels of the respective proteins from the gel on the left. The cytoplasmic domain of DivJ is capable of autophosphorylation and phosphor-transfer to DivK, but not a mutant of DivK that lacks its phospho-acceptor amino acid residue (D53).

## Movie Captions

**Movie S1**

Three dimensional widefield fluorescence images of condensates formed by eYFP fusion of the SpmX-IDR. 5 µM eYFP-tagged SpmX-IDR was incubated in a physiological buffer (50 mM HEPES-KOH, pH 7.4, 0.1 M KCl) for 30 mins followed by microscopy. Slices from 0-0.8 µm show the presence of collapsed eYFP-IDR condensates on glass surface treated with aminosilane. Slices above 1 µm reveal the presence of spherical condensates diffusing in the buffer.

**Movie S2**

Sequential two-color, three dimensional, widefield fluorescence imaging of condensates formed by eYFP-SpmX-IDR and DivJ. 5 µM eYFP-tagged SpmX-IDR was incubated in the presence of 5 µM DivJ (ΔTM, 1% labeled with JF646) in a physiological buffer (50 mM HEPES-KOH, pH 7.4, 0.1 M KCl) for 30 mins followed by microscopy. The SpmX and DivJ channels are separated in time by 8 ms. DivJ is enriched within SpmX-IDR condensates *in vitro*.

**Movie S3.**

3 dimensional confocal imaging of PopZ condensates at room temperature. 5 µM PopZ (1% labeled with Atto488) was reconstituted in a physiological buffer (50mM HEPES-KOH, pH 7.4, 0.1 M KCl) in an aminosilane treated glass bottom chamber for 30 mins. PopZ exhibited an arrested, beads on a string-like morphology that formed an extended network of condensates in 3 dimensions. The 3D stack was acquired using a 63x oil objective with 280 nm slices.

**Movie S4.**

3 dimensional images of PopZ were acquired as in Movie S3 followed by deconvolution in Fiji. A 12 µM by 12 µM by 6 µM volume was selected from a Z-stack and exported into Imaris. A Gaussian filter spanning 1 pixel was applied to the image and 3D reconstruction was performed. Various perspectives of the condensates were then exported into a movie.

**Movie S5.**

Observation of dynamic hollow condensates of 5 µM SpmX (ΔTM, 5% labeled with Cy3) incubated in the presence of 1 mM ATP in a physiological buffer (50 mM HEPES-KOH, pH 7.4, 0.1 M KCl, 5 mM MgCl_2_) at room temperature for 30 minutes. Images were captured every 10 seconds and the movie was recorded for 25 minutes. The jitter in the hollow condensates is due to bubble fusion events along the z-axis (into the plane of the image).

## Materials and Methods

### 1. Strain Engineering

**Table S1.**
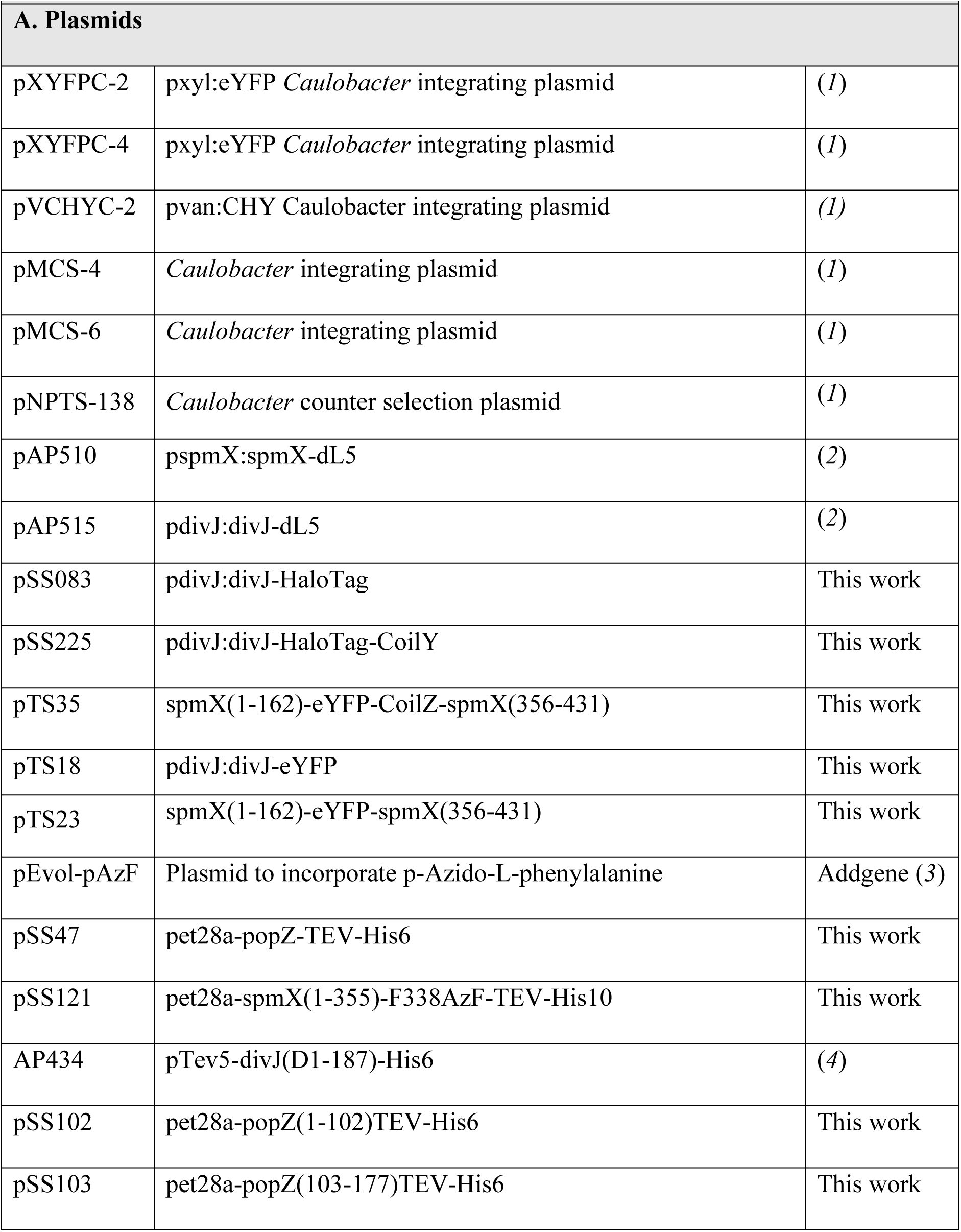
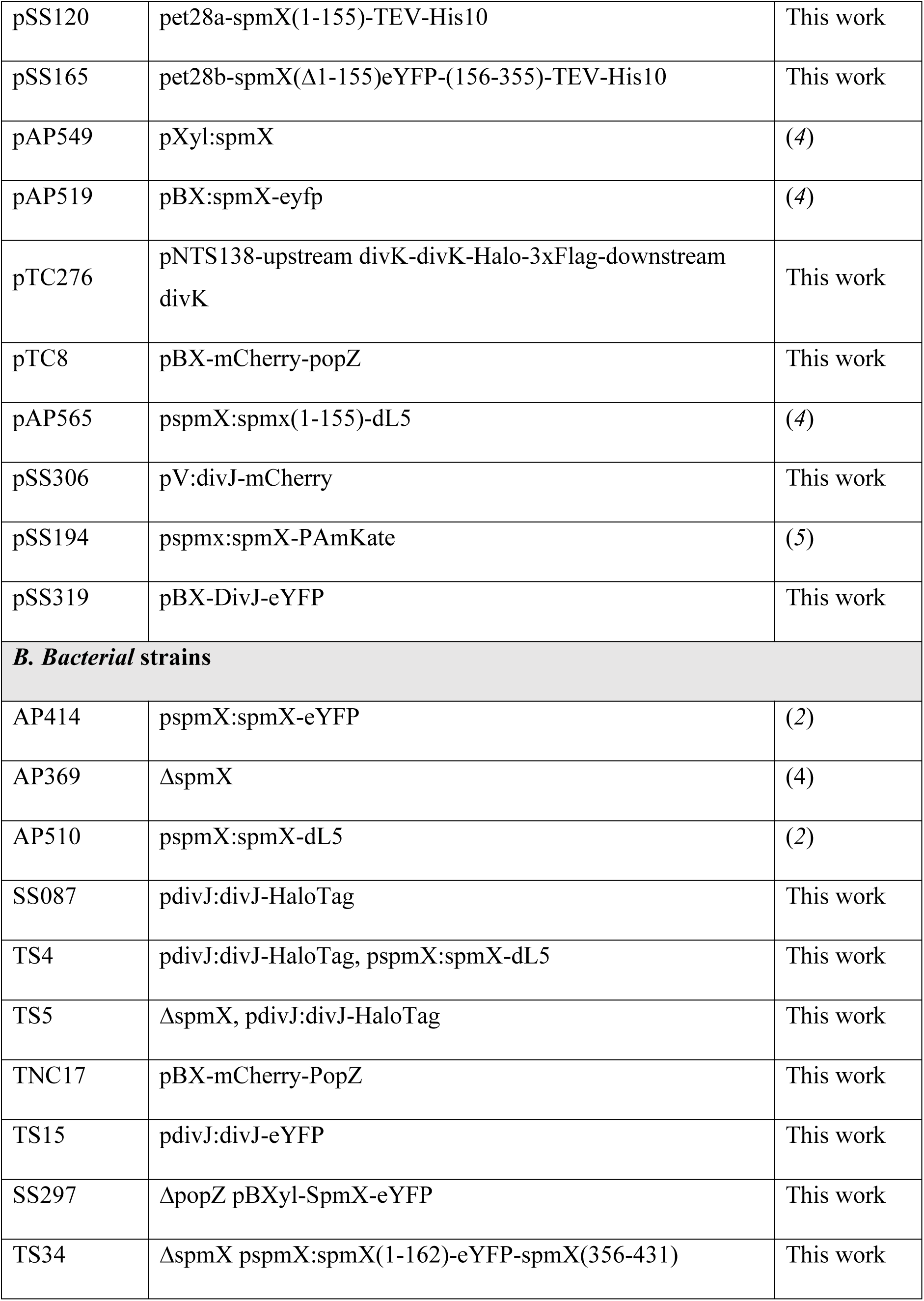
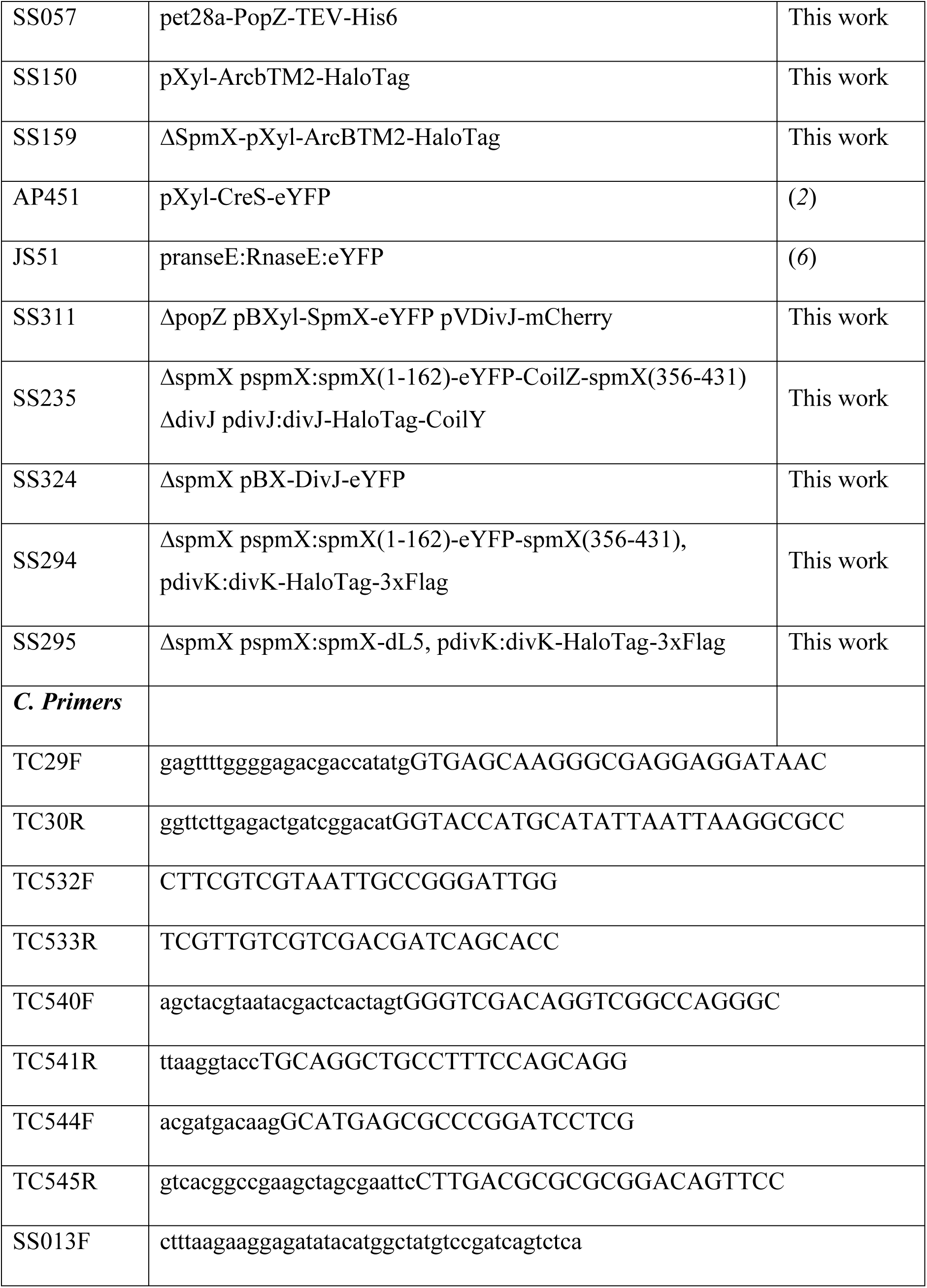
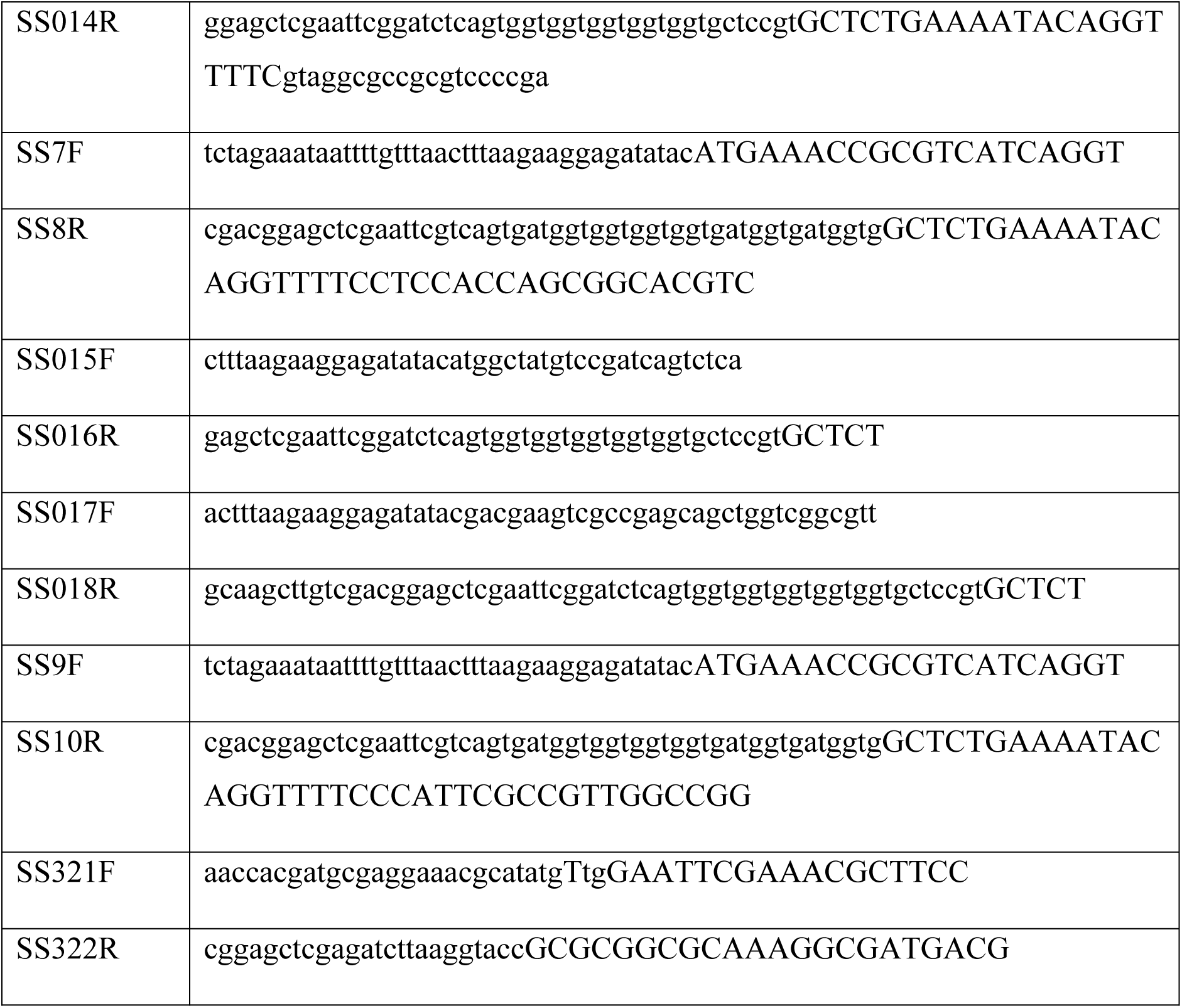
Plasmids and strains used in the study.

#### 1.1 Construction of strains

pTS18 was constructed by inserting eYFP into pAP515 using KpnI and NheI restriction sites by T4 ligation. pTS23 was constructed by spmX(1-162)-eYFP-spmX(356-431) synthesis and insertion into plasmid pMCS-4 using EcoRI and NheI restriction sites by Synbio technologies. pTC276 was constructed by amplification of divK plus 495 base pairs upstream of divK with primers TC540F and TC541R and 800 base pairs downstream of divK with primers TC544F and TC545R from NA1000 genomic DNA. These two DNA fragments along with a gBlock containing the sequence for Halo-3xFlag synthesized by IDT with 5’ and 3’ overlapping sequences were inserted into the plasmid pNPTS-138 using restriction sites SpeI and EcoRI by Gibson assembly. pTC8 was constructed by amplification of mCherry with primers TC29F and TC30R and inserted into a pBX-popZ plasmid (*7*) using the NdeI restriction site and Gibson assembly. pSS083 was constructed by inserting a synthesized piece of HaloTag with appropriate overhangs into pAP515 using KpnI and NheI restriction sites by T4 ligation. pSS225 was constructed by inserting a synthesized piece of HaloTag-CoilY with appropriate overhangs into pAP515 using KpnI and NheI restriction sites by T4 ligation. pTS35 was constructed by inserting a synthesized fragment of spmX(1-162)-eYFP-CoilZ-spmX(356-431), into pTS23 using NheI and EcoRI restriction sites by T4 ligation. pSS47 was constructed by amplifying PopZ from *Caulobacter* genomic DNA using primer SS013F and adding the TEV and His6 sequence on the reverse primer SS014R. The resulting fragment was inserted into pet28a vector digested using EcoRI restriction site by T4 ligation.

The Amber codon containing SpmX(1-431) was synthesized and inserted into a pet28a vector. pSS121 was constructed by amplifying the sequence for spmX(1-355) using primer SS7F and adding the TEV site and His10 on the reverse primer SS8R. The resulting fragment was inserted into pet28a vector digested using NcoI and BamHI restriction sites. pSS102 was constructed by amplifying PopZ(1-102) with TEV and His6 from pSS47 using primer SS15F and SS16R followed by insertion into pet28a vector digested using EcoRI restriction site by T4 ligation. pSS103 was constructed by amplifying PopZ(103-177) with TEV and His6 from pSS47 using primer SS17F and SS18R followed by insertion into pet28a vector digested using EcoRI restriction site by T4 ligation. pSS120 was constructed by amplifying SpmX(1-155) from pSS43 using primers SS9F and SS10R and inserting the resulting fragment into pet28a vector digested using NcoI and BamHI restrictions sites. pSS165 was constructed by synthesizing an eYFP sequence fused to SpmX(156-355)-TEV-His10 sequence and inserting the resulting fragment into pet28a vector digested using NcoI and BamHI restrictions enzymes. pSS306 was constructed by amplifying the DivJ coding sequence from pAP515 using primers SS321F and SS322R and inserting the resulting fragment into pVCHYC-2 digested using NdeI and KpnI restriction enzymes. pSS319 was constructed by amplifying the DivJ-eYFP sequence from pTS18 and inserting the resulting fragment into pBXMCS-4 plasmid digested using NcoI and KpnI.

TS4 was constructed by transduction of SS087 with fCr30 phage carrying pspmX:spmX-dL5 and selected by gentamycin resistance. TS5 was constructed by transduction of *DspmX* with phage carrying pdivJ:divJ-Halo and selected by chloramphenicol resistance. TNC17 was constructed by electroporating NA1000 cells with plasmid pTC8 and selection for gentamycin resistance. TS15 was constructed by electroporation of *DdivJ* cells with plasmid pTS18 and selection for chloramphenicol resistance. TS34 was constructed by electroporation of SS087 cells with plasmid pTS23 and selection for gentamycin resistance. SS057 was constructed by electroporating pSS47 in BL21(DE3) cells. SS311 was constructed by electroporating pSS306 in SS297. SS235 was constructed by electroporating pTS35 into AP369 cells followed by a second electroporation with the plasmid pSS225. SS324 was constructed by electroporating pSS319 into AP369.

SS294 and SS295 were constructed by electroporating TS4 or TS34 with plasmid pTC276. Following selection for kanamycin resistance, recombination of transgenic cells was allowed to occur overnight in PYE liquid culture containing no antibiotics. Recombinant lines having lost the *sacB* gene in the pNPTS-138 backbone which confers sucrose sensitivity were selected for on PYE plates containing 3% sucrose. Recombinant lines containing divK-Halo-3xFlag were confirmed by PCR of lysed cells using primers TC532F and TC533R. The DNA for the transmembrane domain of the protein ArcB (Aerobic respiration control sensor protein, P0AEC3) from *E. coli* was synthesized with a Glycine-Serine linker (GSx4) with overhangs to complement the Xylose locus on one end and the HaloTag locus on the other. The plasmid was then constructed via Gibson assembly into the vector backbone pBXMCS-6 (*8*). The resulting plasmid was electroporated into WT or Δ*spmX* strains and selected with appropriate antibiotics to obtain SS150 and SS159, respectively. The peptide sequence of the transmembrane domain is : MKQIRLLAQYYVDLMMKLGLVRFSMLLALALVVLAIVVQMAVTMVLHGQVESIDVIRSIFFGLLITPWAVYFLSVVVEQLEESRQR.

### 2. Recombinant protein purification and labeling

#### 2.1. Protein expression

All recombinant proteins used in this study were purified using IPTG inducible T7 expression vectors. The cytoplasmic portion of SpmX(amino acid (AA) residues 1-355) was expressed in a modified pet28a vector to contain a C-terminal TEV proteolytic site followed by a polyhistidine tag (His-10). The codon encoding for residue 338 was modified from a codon for phenylalanine to an Amber codon. This was done to enable labeling the protein molecule with a single dye using click chemistry. BL21(DE3) cells were co-transformed with the protein bearing pet28a-SpmX(1-355)-TEV-His10 plasmid and the tRNA synthetase and tRNA expressing plasmid pEvol-pAzF (*3*) for incorporation of p-Azido-L-phenylalanine, into proteins in response to the amber codon, TAG. pEVOL-pAzF was a gift from Dr. Peter Schultz (Addgene plasmid # 31186). BL21 cells containing these plasmids were grown overnight in 2x-YT media at 37°C. Next morning, cells were diluted 100-fold in 1 L of 2x-YT media supplemented with Kanamycin (20 µg/mL) and Chloramphenicol (30 µg/mL) and grown in 2.8 L baffled flasks at 37 °C until they reached an optical density at 600 nm (OD-600) of 0.3. At this stage, 238 mg of IPTG, 200 mg of L-Arabinose and 200 mg of p-Azido-L-Phenylalanine (dissolved in 2 mL of 0.1 N NaOH) were added to the flasks. The flasks were then moved to a 30 °C incubator where protein induction was carried out for 4-5 hours. After induction, the cells were washed using cell wash buffer (50 mM Tris, pH 7.4, 500 mM NaCl) and pelleted in 50 mL conical tubes by spinning at 3700xG for 15 mins. The pellets were saved in a freezer at -80 °C until they were ready to be used. SpmXΔIDR(AA 1-155), PopZ(AA 1-177), PopZ (AA 1-102), PopZ (AA 103-177), DivJΔTM(AA 188-597), DivK, and DivK(D53N) were all expressed without Amber codons in pet28a vector containing TEV protease and polyHistidine sites. We observed that SpmX(156-355) was unstable in *E. coli*. To circumvent this, we expressed SpmX(AA 156-355) as a N-terminal fusion to eYFP. The resulting protein eYFP-SpmX-IDR(AA 156-355)-TEV-His10 was more stable than the untagged SpmX-IDR and could be purified to concentrations up to 25 µM. Proteins expressed without Amber codons were purified using the same protocol as above with appropriate antibiotics, except that the cells were grown in LB and only IPTG was added during the induction step.

#### 2.2. Protein purification

For purifying proteins from BL21 cells, frozen pellets were thawed on ice in 50 mL conical flasks with pre-chilled lysis buffer (50 mM HEPES – KOH pH 7.6, 500 mM KCl, Roche Protease Inhibitors tablet (0.5 tablet/L of culture), 25 mM imidazole, 200 U benzonase nuclease (0.3 µL/ 1 L culture). Cells were resuspended in lysis buffer and lysed using sonication. The sonicated sample was centrifuged at 29000XG for 45 mins in a JS-20 rotor at 4 °C. The supernatant was collected and incubated with 2 mL of pre-equilibrated Ni-NTA agarose slurry (50%) for each liter spun down cells. The slurry was nutated for 2 hours at 4 °C. The slurry was then washed three times with Ni-NTA wash buffer (50 mM HEPES-KOH pH 7.6, 500 mM KCl, 25 mM imidazole) and protein was eluted from the slurry by incubating the slurry with elution buffer (50 mM HEPES-KOH pH 7.6, 500 mM KCl, 250 mM imidazole,10% glycerol) for 20 minutes and collecting the flow-through. Proteins were then dialyzed twice against dialysis buffer (50 mM HEPES-KOH pH 7.4, 100 mM KCl, 10% glycerol). DTT was avoided in dialyses where the protein was to be subsequently labeled using a fluorescent dye, as DTT reduced fluorophores and affected multivalency within condensates. Proteins that did not contain an IDR were further purified by size exclusion chromatography. Proteins were concentrated to appropriate stocks and flash frozen in liquid Nitrogen and frozen at -80 °C until needed.

#### 2.3. Protein labeling

For SpmX labeling with Cy3-DBCO using click chemistry, the dye reconstituted in the dialysis buffer and incubated with the eluted protein at 20x molar excess overnight at 4 °C. In the case of proteins that were labeled using Atto-488 NHS dye, labeling was performed by using 20x molar excess of Atto488-NHS for 2 hours at 4 °C. Free dye was removed from the samples by gravity assisted chromatography on a PD-10 column (Cytiva, 17-0851-01). Proteins were concentrated to appropriate stocks and flash frozen in liquid Nitrogen and frozen at -80 °C until needed. Labeled and unlabeled proteins were mixed at the requisite ratio for one hour before the experiments.

### 3. *In vitro* reconstitution

#### 3.1. Imaging chamber preparation

Droplet experiments for SpmX phase separation were performed in glass bottom 384 well plates (MatTek, PBK384G-1.5-C). The wells were treated by flushing them with Nitrogen gas followed by the addition of 50 µL freshly prepared 1 M KOH in milliQ water. The plate was centrifuged for 10 seconds at 600xG to ensure that the liquid has settled down, followed by a 10 minute incubation at room temperature with slow shaking. Next, the wells were washed with 50 µL milliQ water five times. After the final wash, 20 µLof (3-Aminopropyl)triethoxysilane (APTES, Vector labs) was added to each well. The plate was centrifuged for 10 seconds at 600XG to ensure that the liquid has settled down. The wells were covered with an aluminum foil and incubated for 1 hour at room temperature with slow shaking.

The wells were washed five times with 50 µL PBS per wash and five times with 50 µL water per wash. The wells were dried for 1-2 hours followed by imaging. In general, we observed a deterioration in the quality of the chemically modified surface after one day of the treatment. Therefore, this procedure was performed on the day of the experiments.

#### 3.2. Protein treatment

Poly-histidine tags were cleaved from proteins by treating them with 3 units of TEV protease (Sigma) overnight at 4 °C based on the manufacturer’s guidelines. TEV-proteolyzed unlabeled proteins were mixed with TEV-proteolyzed dye-labeled proteins at the indicated mole %, which was 1% for PopZ-Atto488, 5% for SpmX-Cy3 (or SpmXΔIDR-Cy3), 1% for DivJ-Atto488 and 1% for DivJ-Cy3. eYFP fusion of SpmX-IDR was used without any additional mixing steps. PopZ samples were prepared by mixing labeled and unlabeled protein to obtain a stock, followed by heating the samples in PCR tubes at 98 °C for 8 mins.

##### 3.2.1. Condensate imaging

Microscopic observation of condensates was performed by diluting the TEV-proteolyzed proteins to 5 µM in a physiological buffer (50 mM HEPES-KOH pH 7.4, 100 mM KCl). Where needed, proteins were mixed at the respective concentrations in the physiological buffer. Protein solutions were then incubated at room temperature for 30 mins followed by addition of 30 µL samples into the APTES treated chambers in glass bottom 384-well plates. The plate was then mounted on the microscope stage. Condensates could be observed in solution, away from the glass followed by settling to the bottom of the chamber within 10-20 mins. The eYFP-fused SpmX-IDR protein was an exception to this observation. eYFP-IDR condensates collapsed upon touching the APTES treated glass surface. Other surface modifications were tried for eYFP-IDR (KOH, Poly-L-Lysine, BSA, Poly-L-Lysine-PEG, or untreated glass with or without Argon plasma etching), with no success. As a result of this surface induced instability, eYFP-IDR condensates (with or without DivJ-Cy3) were imaged before they fused on the glass surface.

##### 3.2.2. Salt-dependent condensate phase diagram imaging

For imaging condensates without any salt, an aliquot of concentrated protein was dialyzed against a buffer lacking any KCl (50 mM HEPES-KOH pH 7.4, 10% glycerol) using mini-dialysis cassettes (Slide-A-Lyzer™ MINI Dialysis Device, 3.5 K MWCO, 0.1 mL) for 1 hour. The dialyzed and desalted protein was then taken up at appropriate concentrations in the imaging buffer (50 mM HEPES-KOH pH 7.4, 10% glycerol). For 0.1 M KCl, the purified protein was diluted to appropriate concentrations in imaging buffer containing 0.1 M KCl, and for higher KCl concentrations (0.2-0.5 M), additional KCl dissolved in the imaging buffer was supplemented into the reactions.

##### 3.2.3. Condensate imaging under chemical perturbations

For experiments requiring ATP addition, 5 µM protein samples were prepared as above and incubated at room temperature for 30 mins before addition of the requisite concentrations of ATP in imaging buffer (50 mM HEPES-KOH pH 7.4, 100 mM KCl, 5 mM MgCl_2_). ATP stocks were prepared in water at 500 mM concentration, and pH was adjusted to 7.4 by adding 1 N NaOH. Subsequent dilutions of the 500 mM ATP stock were made in the imaging buffer (50 mM HEPES-KOH pH 7.4, 100 mM KCl, 5 mM MgCl_2_). Addition of the highest concentration of ATP led to a change in the ionic strength of the solution from 115 to 123 (7%). Using a control buffer with matched ionic strength (50 mM HEPES-KOH pH 7.4, 108 mM KCl and 5 mM MgCl_2_), we validated that the observed effects were not due to changes in ionic strength. LA stocks were made in DMSO at 500 µM and dilutions of the stock were made in imaging buffer (50 mM HEPES-KOH pH 7.4, 100 mM KCl). As a control for experiment requiring LA addition, an equal volume of DMSO was added to the protein samples. 1,6-HD was purchased as a 99% aqueous solution and was added at desired concentrations to proteins. In this case, an equivalent volume of water was added to the untreated samples as a control.

#### 3.3. Turbidity measurement using absorbance

To assess turbidity in protein samples, absorbance at 600 nm was measured in 384-well plate format using a Varioskan LUX multimode microplate reader (ThermoFisher). Unlabeled SpmX or PopZ were reconstituted in a physiological buffer (50 mM HEPES-KOH pH 7.4, 0.1 M KCl) at increasing concentrations. Protein samples were added to a 384-well polystyrene plate and incubated for 30 mins at room temperature before measuring absorbance on a plate reader. As a control purified DivJ and HaloTag were used at the same concentration range and did not exhibit any concentration dependent turbidity. In all cases, the buffer only background was subtracted from the absorbance values.

### 4. In vitro DivJ kinase assays

#### 4.1. Liposomes

DivJ phosphorylation assay on liposomes was performed on nitrocellulose blots as described previously (*9, 10*). Small unilamellar vesicles (SUVs) were prepared by dissolving polar phospholipids (Avanti Polar Lipids; Alabaster, AL) in chloroform at a 9:1 molar ratio (9 parts 1,2-dioleoyl-sn-glycero-3-phospho-(1′-rac-glycerol; sodium salt; di-oleoyl-phosphatidyl glycerol: product 840475) to 1 part 1,2-dioleoyl-sn-glycero-3-[(N-(5-amino-1-carboxypentyl)iminodiacetic acid)succinyl] (nickel salt; DGS-NTA(Ni): product 790404C) in a glass scintillation vial. The chloroform solvent was removed under a slow and steady flow of Nitrogen gas for 2 hours to obtain a dry, thin film in the vial. The film was rehydrated in the kinase buffer (25 mM HEPES-KOH, pH 7.4, 50 mM KCl, 25 mM NaCl, 5 mM MgCl_2_) to yield 25 mg/mL liposome sample. The buffer was vortexed with the film for 10 min until the lipid was fully dissolved. The aqueous lipid mixture was then subjected to 10 freeze/thaw cycles in liquid nitrogen and a 37 °C water bath. The freeze-thaw mixture was then extruded for 11 passes through 100 nm pores of a polycarbonate filter using the Avanti Mini-Extruder. The resulting SUVs were collected in a glass vial and stored at 4 °C for up to 1 week. 5 µM DivJ-His_6_ was incubated in kinase buffer for 5 mins at room temperature. Following this, serial dilutions of SUV stock solutions were added such that the volume of liposome solution added in each case was the same. As a control SUVs prepared without DGS-NTA lipids were also incubated with DivJ at multiple densities. The incubations were carried out in PCR tubes for 45 mins at room temperature. 1 μCi [γ−^32^P] ATP was added to each reaction tube and the tubes were incubated for 2 mins at room temperature. Each reaction was performed in triplicate. 4 μL of each reaction was spotted on the nitrocellulose membrane and dried under a lamp for 40 mins. Following the drying, the membrane was washed five times with 50 mL of 12.5 mM sodium pyrophosphate buffer, pH 10. The membrane was dried for 1 hour under an incandescent lamp, wrapped in a plastic wrap and transferred to a phosphor imaging screen box. The screen was exposed to the membrane for 3 hours followed by phosphorimaging on a Typhoon imager. The estimation of number of DivJ molecules per liposome is based on similar experiments with the protein kinase CckA (*9*). The goal of this estimation was to assess whether the number of DivJ molecules per liposome is limited by geometrical considerations (area) or concentration of protein and Nickel-NTA sites. To estimate the number of available binding sites for DivJ molecules on a liposome, we calculated the total surface area of the liposome. DOPG liposomes extruded through 100 nm pores have vesicle diameters of 97-106 nm (*11*), that we approximated as 100 nm for simplification of calculation. Based on these calculations, we estimated that at an equal mass of lipid to DivJ protein, the density of DivJ was ∼1100 molecules per liposome. The data from spot assays in triplicate was averaged and fit using the Hill-Langmuir equation:

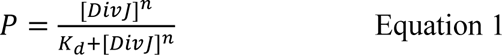

In equation 1, *P* denotes the fraction of DivJ phosphorylated, denoted by the end point kinase activity, [*DivJ*] denotes the estimated DivJ density, *K_d_* is the apparent dissociation constant and *n* is the Hill coefficient.

#### 4.2. In vitro phosphor-transfer to DivK

To test DivJ’s autokinase activity and its phosphotransfer to DivK, we performed assays in a kinase buffer (50 mM HEPES-KOH pH 7.4, 25 mM NaCl, 25 mM KCl, 5 mM MgCl_2_) with equimolar protein concentrations as needed. 5 µM DivJ, 5 µM DivJ and 5 µM DivK, or 5 µM DivJ and 5 µM DivK(D53N) were incubated in the kinase buffer for 30 mins at room temperature. The kinase reaction was carried out by the addition of 1 μCi [γ−^32^P] ATP for 5 mins at room temperature. The reactions were quenched by adding 4% SDS followed by separation of proteins by SDS-PAGE in duplicate. One part of the gel was covered in a plastic wrap and transferred to a phosphor imaging screen box. The screen was exposed to the gel for 3 hours followed by phosphor imaging on a Typhoon imager. The other part of the gel was stained using silver chloride and imaged on a protein gel imager.

### 5. Caulobacter protocols

Unless and otherwise stated, *Caulobacter* cells were grown from frozen stocks in PYE overnight with appropriate antibiotics. The following day, cells were diluted in M2G to OD-600 0.01 and grown to OD-600 0.3-0.4. Imaging was always performed on agarose pads (composed of 1.5% (w/w) of low melting point agarose (Invitrogen) in M2G).

#### 5.1. Imaging

For imaging experiments requiring dye labeling, cells growing in M2G in exponential phase (OD 0.3) were washed with M2G by centrifugation, then labeled with 5 nM (final concentration) JF549-HaloTag (*12*) by adding a 500 nM stock (in DMSO) solution to the cells. For SpmX-dL5 imaging, cells were labeled with 10 nM (final concentration) of the dye MG-ester (1 µM stock in Ethanol, 5% Acetic acid) (*13*). The cells were incubated with the dye and shaken for 20 minutes at room temperature in the dark, then washed by centrifugation (3 min, 8000 RPM, 28 °C (Eppendorf 5430R)) and resuspension in M2G three times. After the final wash, cells were resuspended in ∼20-50 μL of M2G, producing a concentrated cell suspension. To this suspension, ∼1 nM of fiducial markers was added (Molecular Probes, 540/560 carboxylate-modified FluoSpheres, 100 nm diameter). Next, 1-2 μL of the cell/fiducial mixture was deposited onto an agarose pad, mounted onto an argon plasma-etched glass slide (22×22 mm No. 1.5, Fisher), covered with a smaller (18×18 mm, No. 1, Fisher) plasma-etched slide, sealed with wax, and imaged immediately. For imaging cells with fluorescent proteins, a similar protocol was followed without the dye labeling steps and washing. Cells growing in exponential phase were washed once with M2G, then resuspended in ∼20-50 μL of M2G. 1-2 μL of this mixture was deposited onto an agarose pad, mounted onto a plasma-etched glass slide, and imaged immediately. For confocal imaging of SpmX-dL5 samples, all steps were carried as above except that no fiduciary markers were used for imaging.

#### 5.2. In vivo phosphorylation

##### 5.2.1. DivJ phosphorylation measurements

In vivo phosphorylation measurements were carried out as described previously (*14*) with the following modifications. Cells from frozen stocks were inoculated into M5G low phosphate medium and grown overnight at 30 °C until the OD-600 was between 0.35-0.4. Cultures were normalized to the lowest OD culture in the batch. 1 mL of cells from each culture pulsed with 1 μCi [γ^32^P]-ATP having a specific activity of 30 Ci/mmol (Perkin Elmer) for 5 minutes at room temperature. Immunoprecipitations were performed using Protein A agarose beads (Roche). DivJ was immunoprecipitated using an anti-DivJ antiserum (*15*). The immunoprecipitated sample was run on two SDS-PAGE gels. One of the gels was covered in a plastic wrap, dried, and transferred to a phosphor imaging screen box. The screen was exposed to the gel for 5 days followed by phosphor imaging on a Typhoon imager. Proteins separated on the second gel were transferred to a PVDF membrane and DivJ levels were measured using Western blots by probing the protein with anti-HaloTag antibody (Promega). Due to the requirement of growing cells in low phosphate media, this assay could not be used to measure phosphorylated DivJ levels under varying glucose conditions.

##### 5.2.2. DivK phosphorylation measurements

To measure phosphorylated DivK levels as a function of available glucose, cells were grown in M2 minimal media supplemented with 1 mM, 0.5 mM or 0.2 mM glucose at 30 °C. 1 mL cells were harvested at an OD-600 of 0.35-0.4. Harvested cells were normalized to the lowest OD-600 of the collected cells and pelleted by centrifugation at 20000XG for 2 mins and resuspended in 75 µL lysis buffer (10 mM Tris-HCl, pH 7.0, 4% SDS, 5 U DNaseI (ThermoFisher), 0.2 tablets of PhosStop (Roche)). Cell pellets were lysed for 5 mins at room temperature. One of the cell pellet samples was lysed at 98 °C for 5 min to degrade phosphorylated DivK and serve as a negative control in the assay. After lysis, the samples were spun at 16000xG for 5 mins at room temperature. 12 µL of the supernatant was carefully pipetted from the tubes and mixed with 12 µL of 2x sample buffer (20 mM Tris-HCl, pH 7.0, 6% SDS, 10 mM EDTA, 20% glycerol (v/v), and 10% β-mercaptoethanol (v/v)). 20 µL of this sample was loaded on the PhosTag acrylamide gels (Fujifilm Wako chemicals) for separation. Phos-tag SDS-PAGE gels were prepared with 50 μM Phos-tag acrylamide and 100 μM ZnCl_2_ and run at 4 °C at 80 V. Prior to transfer by western blot, gels were washed three times for 10 minutes in transfer buffer supplemented with 10 mM EDTA at room temperature to remove Zn^2+^ from the gel and washed once with transfer buffer without EDTA. Proteins were transferred using a semi-dry platform to PVDF membranes at 25 V for 2 hours at room temperature. DivK protein separated based on its phosphorylation state was transferred to a PVDF membrane and detected using a chemiluminescent substrate (SuperSignal West PICO PLUS, Thermo Pierce) after incubation with mouse anti-Flag (1:10000) primary antibody (Sigma, F1804) and a polyclonal goat anti-mouse HRP (1:10000) conjugated secondary antibody (Abcam). Phosphorylated and non-phosphorylated forms of DivK were estimated using Fiji from three independent biological replicates.

#### 5.3. In vivo chemical perturbations

To test the effect of different chemical perturbations on fluorescently tagged proteins in vivo, *Caulobacter* cells were grown from frozen stocks in PYE to an OD-600 of 0.3-0.4 (or 0.2 for *ΔpopZ* strains). Protein production was induced for four hours by supplementing the growth media with 0.3% Xylose. After induction, 1 mL cells were spun down (12000xG for 2 mins) and resuspended in M2G (for CCCP treatment) or M2 minimal media (for LA or 1,6-HD treatment). 100 µM CCCP (10 mins), 5 µM LA (30 mins), or 5% 1,6-HD (30 mins) were added to the resuspended cultures followed by shaking at 30 °C for the indicated amount of time. After the treatment, 1-2 μL of cell samples were deposited onto an agarose pad, mounted onto a glass coverslip, and imaged immediately. Ten, 80 µm fields of view were imaged per sample, and this routine was repeated for three biological replicates. For measuring cell health under solute treatment, a cell impermeant fluorescent dye (Live-or-Dye 640-662, Biotium) was used. Lyophilized dye powder was reconstituted in 50 µL anhydrous DMSO based on the manufacturer’s protocols. 1 µL of the dye stock was added to 1 mL cells treated with various perturbations, and the samples were incubated on a rotatory shaker at 30 °C for 30 mins. As a control an equivalent volume of DMSO was added to untreated cells. Cells were washed thrice with 1 mL M2G media followed by imaging on agarose pads. Cells that sequestered the dye within them were at least five-fold brighter than the healthy cells. Accordingly, a threshold was applied to obtain the number of cells with or without membrane leakage using bespoke code in Matlab.

#### 5.4. Glucose dependent measurements

##### 5.4.1. Cell length

For cell length measurements, cells cultures were started from frozen stocks in PYE media. These cultures were collected in exponential phase and washed once with M2 minimal media without Glucose. Cells were diluted to OD-600 0.05 in respective media (M2G (1 mM), M2G (0.5 mM) etc.) in triplicate and grown to exponential phase (OD-600 of 0.2-0.4). These cultures were diluted 10-fold and imaged on M2G-agarose pads immediately. OD-600 for each culture was measured, 1 mL of each culture was centrifuged at 16000xG for 2 mins and the media was aspirated. These pellets were frozen in liquid nitrogen and were stored at -80 °C prior to ATP measurements. For intracellular ATP measurements, the pellets were thawed and resuspended in 300 µL of M2 minimal media and the assay was performed as per the manufacturer’s protocols (BacTiter-Glo, Promega) (*16*). For each sample, luminescence measurements were made in 3 wells and average value was taken from 9 wells per strain and condition (For example: WT strain in M2G had pellets from 3 biological replicates, each pellet was resuspended in 300 µL lysis buffer and was split into 3 samples of 100 µL per sample for the luminescence assay). Cell length analysis was performed using MicrobeJ plugin for Fiji (*17, 18*). We employed manual mode detection for long and filamentous cells (representing SpmXΔIDR, ΔSpmX or DivJ(H338A) phenotypes) that could not be selected by the automated detection using a reasonable set of thresholding parameters.

##### 5.4.2 Growth curve

For growth curve measurements, cells in exponential phase were back diluted to O.D. 0.02 and grown in 96 well plates at 28 °C with constant shaking in between measurements on a Tecan Safire multi-well plate reader. OD-600 was measured every 10 minutes.

#### 5.5. Western blots

To assess the levels of SpmX in different strains (Fig. S2I), cells were grown in M2 minimal media supplemented with 1 mM glucose at 30 °C. Cells were incubated with or without indicated concentrations of Xylose for 30 mins to enable protein induction. 1 mL cells were harvested at an OD-600 of 0.35-0.4. Harvested cells were normalized to the lowest OD-600 of the collected cells and pelleted by centrifugation at 20000XG for 2 mins and resuspended in 30 µL of 2x sample buffer (20 mM Tris-HCl, pH 7.0, 6% SDS, 10 mM EDTA, 20% glycerol (v/v), and 10% β-mercaptoethanol (v/v)) and incubated at 98 °C for 10 mins. 20 µL of this sample was loaded on Mini-PROTEAN® Precast Mini PAGE Gels (Biorad) for separation. Proteins separated via SDS-PAGE were transferred using a semi-dry platform to PVDF membranes at 80 V for 1 hour at room temperature. SpmX levels on the PVDF membrane were detected using a chemiluminescent substrate (SuperSignal West PICO PLUS, Thermo Pierce) after incubation with an anti-SpmX (1:10000) primary antibody raised against the SpmX-IDR (*19*) and a polyclonal goat anti-mouse HRP (1:10000) conjugated secondary antibody (Abcam). SpmX levels were estimated using Fiji from three independent biological replicates.

### 6. Microscopy

#### 6.1. Diffraction-limited wide field imaging (phase contrast and fluorescence)

6.1.1. Data acquisition: Cells immobilized on agarose pads or *in vitro* samples in multi-well glass bottom plates were imaged on a LED based (Lumencor, SpectraX) multi-color epifluorescence microscope consisting of a Leica Dmi8 stand equipped with an immersion oil phase contrast objective (100x, HC PL APO, 1.4NA) and an EMCCD camera (Hamamatsu, C9100 02 CL).

**Table S2.**
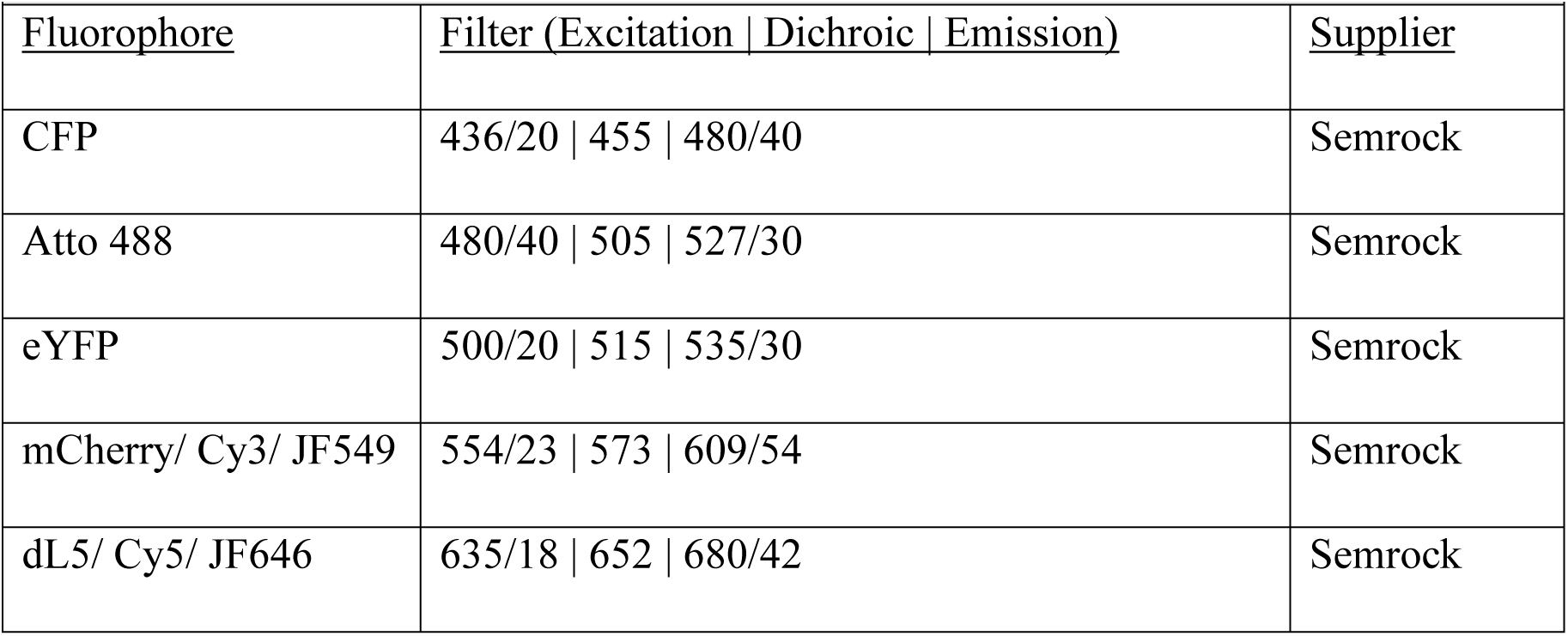
Filters used.

6.1.2. Analyses for condensate intensity, either at a single time point or as a time series were performed using bespoke software written in Matlab. Cell length analyses were performed using MicrobeJ (*17*). Analyses of fluorescent clusters in live cells was performed using bespoke software in Matlab. A binary mask was generated using the phase contrast images of the cells. In each cell within the mask, a threshold was applied to obtain all clusters within a cell. The mean fluorescence intensity from a cellular region not containing the clusters was subtracted from the mean fluorescence intensity of the clusters. This number was divided by the mean fluorescence intensity of the cell for normalization. The automated threshold did not work robustly for RnaseE-eYFP and DivJ-eYFP cells under conditions where the clusters intensities were very low. In these cases, a manual step was added on top of the first threshold step. For 2-color analyses (Fig. S4A), a binary mask was created using the phase contrast images followed by thresholding the SpmX channel to detect and quantify fluorescent clusters. The thresholded image for SpmX channel was then converted to a binary mask and applied to the DivJ channel. Fluorescence value calculations and normalization were performed as above.

#### 6.2. Confocal imaging and FRAP

##### 6.2.1. Data acquisition

Confocal microscopy was employed for imaging *in vitro* condensates and *E. coli* cells. Samples were imaged at room temperature on a spinning disk confocal microscope (Leica DMI6000B custom-adapted with a Yokogawa CSU-X1 spinning disk head; a Photometrics Evolve 512 EMCCD camera; and Intelligent Imaging Innovations SlideBook software, Vector FRAP, LaserStack), with a 100x oil objective (HCX Pl APO, 1.4 NA; Leica). eYFP fluorescence was imaged by excitation at 514 nm and emission with a YFP 540/15 filter (Semrock) and 445/514/561 nm Yokogawa dichroic beamsplitter (Semrock) under the following conditions: 100 ms exposure every 1 s for 100 time points, with < 2 mW laser power (measured at the fiber), with Adaptive Focus Control active during the entire acquisition. Vector was used to direct the 514 nm laser at full power for photobleaching (∼18 mW, measured at the Vector fiber), which took place between the third and fourth image captures on a region of interest positioned at the center of each selected condensate. Line mode of photobleaching was also used on *Caulobacter* and small condensates.

##### 6.2.2. Analysis

FRAP images were exported as tiff files into a Matlab workspace. Image background was measured from regions that did not have any condensates, averaged, and subtracted from the image. FRAP analyses protocol was developed based on reports elsewhere (*20*). Three regions of interest were selected using the *roipoly* function in Matlab. Region 1 was the bleached region within a condensate, region 2 was a control region on another condensate that was not bleached, and region 3 was the entire bleached condensate. Intensity as a function of time was measured from all the three regions. Intensity from region 2 was subtracted from region 1 and the resulting intensity was divided by the intensity from region 3 to normalize for laser / sample fluctuations. Finally, the resulting signal was normalized such that the pre-bleach signal was set to 1 and the resulting FRAP curves were reported as fractional recovery curves.

#### 6.3. Single-molecule imaging

##### 6.3.1. Single-molecule imaging and data acquisition

Cells immobilized on agarose pads were imaged on the two-color home-built epifluorescence microscope described previously (*21*). Two-dimensional white light transmission images of the cells were recorded before imaging. Fluorescence emission from the sample was collected through a high NA oil-immersion objective (Olympus UPlanSApo 60x/1.40 NA), then filtered by a multi-pass dichroic mirror (Chroma, zt440/514/561/642rpc). For detecting eYFP, a 514 nm long pass filter (Semrock LP02-514RE) and a bandpass filter (Semrock FF01-523/610) were used. For detecting JF549, a 560 nm dichroic beamsplitter (Semrock FF560-FDi01), a 561 nm notch filter (Semrock NF03-561E), and a bandpass filter (Semrock FF01-523/610) were used. eYFP was pumped with 514 nm excitation (MPB Communications fiber laser 514 nm 1 W, ∼0.01 – 0.1 kW/cm^2^). JF549 was excited with a 561 nm laser (Coherent Sapphire 561 100 CW, ∼0.75 kW/cm^2^), and reactivated with a 405 nm laser (Coherent Obis, ∼0.1-10 W/cm^2^). Integration times were 20 ms.

##### 6.3.2. Single-molecule data fitting

Three-dimensional single-molecule data were fit using Easy-DHPSF v2.0, freely available at https://sourceforge.net/projects/easy-dhpsf/ (*22*). Calibration scans were generated by axially scanning a fluorescent bead (540/560 100 nm Fluospheres, Life Technologies) using a piezoelectric stage scanner (Physik Instrumente P-545). This axial calibration scan was used to relate angular lobe orientation to axial position, and to generate templates needed to locate candidate single molecules in the data. Background photons (∼7 ± 1 photons/pixel (mean ± S.D.)) were estimated using a temporal median filter. On average, we detected 958 ± 354 photons (mean ± S.D.) per 20 ms frame for the DivJ-Halo-JF549 molecules (561 nm laser intensity of ∼0.75 kW/cm^2^). Localization precision was estimated with an empirical formula derived from repeatedly localizing single beads under variable background conditions (*21*). Localizations used for SPT analysis had typical localization precisions of 30.4 ± 7.5 nm, 31.5 ± 8.0 nm, and 46.9 ± 11.7 nm in x, y, and z, respectively (mean ± S.D.). Systematic errors from sample drift resulting from mechanical and thermal fluctuations were accounted for by adding ∼1 nM concentration of 540/560, 100 nm Fluospheres (Life Technologies) to the sample and using these as fiducial beads, whose motion was removed from the single-molecule localization data.

##### 6.3.3. Diffusion analysis

Single-molecule trajectories were constructed by connecting SM localizations from consecutive frames. Only trajectories of at least 9 steps (10 frames or 200 ms) were used for MSD analyses. The maximum allowed displacements of DivJ-Halo-JF549 molecules over 20 ms were set to 750 nm. For DivJ-Halo-JF549, we detected a median trajectory length of 17 frames (0.34 s). Diffusion coefficients were calculated from ensemble-averaged 3D MSDs (Equation 1.1) (*23–25*) of individual DivJ molecules using only the first two time lags to avoid long-time effects of confinement and non-Brownian motion. The reported diffusion coefficient errors are diffusion coefficient standard deviations estimated by bootstrapping (50% of the trajectories sampled with replacement 500 times).

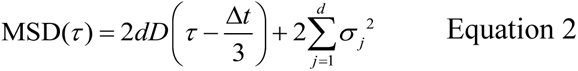

In Equation 2 above, τ is time lag, *Δt* is camera exposure time, *d* is the number of dimensions (3 in this case), *D* is the diffusion coefficient, and *σ_j_* is the localization error in dimension *j*.

To distinguish between diffusion behaviors for polar trajectories, a cumulative distribution function of all displacements in 20ms intervals was calculated (*26*). Since the jumps within this time lag were less than three times the localization precision of the measurements, this procedure could not be used for extracting the diffusion coefficients.

#### 6.4. Airy scan confocal microscopy

##### 6.4.1. Data acquisition

Two color super-resolution imaging of condensates or live *Caulobacter* cells was performed on an inverted laser scanning confocal microscope (Zeiss LSM880) equipped with an AiryScan module (*27*). AiryScan alignment was performed as per the Zeiss manual in both channels. Condensate imaging (Fig. 2F) was performed using a PLAN APO 63x, 1.4 NA oil immersion objective. Atto488 was excited using a 488 nm laser (at 2%) and the resulting fluorescence was collected within a detection window spanning 509-541 nm. Cy3 was excited using a 561 nm laser (at 7 %) and the resulting fluorescence was collected within a detection window spanning 570-610 nm. Live cell imaging (Fig. 2G) was performed using an EC PLAN NEO 40x, 1.3 NA oil immersion objective. mCherry was excited using a 561 nm laser (at 1.9%) and the resulting fluorescence was collected within a detection window spanning 570-620 nm. dL5 was excited using a 633 nm laser (at 0.5%) and the fluorescence was collected using a 645 nm long pass filter.

##### 6.4.2. Analyses

Analyses of condensate demixing in vitro was performed using bespoke software written in Matlab. For each image, a binary mask was generated by applying an intensity threshold on the PopZ channel. The mask was applied on the SpmX channel followed by the application of a spot detection algorithm to count the number of SpmX clusters. A similar analysis was performed for live cell images whereby PopZ assemblies above a set intensity threshold and larger than 1 µm were used to create a binary mask. This mask was applied on the SpmX channel followed by the spot detection analyses as in the previous case, to count the number of SpmX clusters within PopZ microdomain.

#### 6.5. Correlative cryogenic electron microscopy

Cryogenic single molecule imaging of SpmX-PAmKate fusions expressed in live *Caulobacter* cells was performed according to protocols reported previously (*5*). SpmX localizations from four cells were pooled and overlaid on a representative image of *Caulobacter* obtained from cryo-electron tomography. Registration between single molecule fluorescence and cryo-EM images was performed as reported previously (*28*).

## Acknowledgements

pEVOL-pAzF was a gift from Peter Schultz (Addgene plasmid # 31186 ; http://n2t.net/addgene:31186 ; RRID:Addgene_31186)

## References

1. S. F. Banani, H. O. Lee, A. A. Hyman, M. K. Rosen, Biomolecular condensates: organizers of cellular biochemistry. Nature reviews. Molecular cell biology 18, 285–298 (2017).

2. Y. Shin, C. P. Brangwynne, Liquid phase condensation in cell physiology and disease. Science (New York, N.Y.) 357, eaaf4382 (2017).

3. A. Patel et al., ATP as a biological hydrotrope. Science (New York, N.Y.) 356, 753–756 (2017).

4. S. K. Govers, C. Jacobs-Wagner, *Caulobacter crescentus*: model system extraordinaire. Current Biology 30, R1151–R1158 (2020).

5. G. R. Bowman et al., Oligomerization and higher-order assembly contribute to sub- cellular localization of a bacterial scaffold. Molecular Microbiology 90, 776–795 (2013).

6. S. K. Radhakrishnan, M. Thanbichler, P. H. Viollier, The dynamic interplay between a cell fate determinant and a lysozyme homolog drives the asymmetric division cycle of *Caulobacter crescentus*. Genes & development 22, 212–225 (2008).

7. A. M. Perez et al., A localized complex of two protein oligomers controls the orientation of cell polarity. mBio 8, 2238 (2017).

8. A. Gahlmann et al., Quantitative multicolor subdiffraction imaging of bacterial protein ultrastructures in three dimensions. Nano Letters 13, 987–993 (2013).

9. J. B. Grimm et al., A general method to improve fluorophores for live-cell and single-molecule microscopy. Nature methods 12, 244–250 (2015).

10. L. Weimann et al., A quantitative comparison of single-dye tracking analysis tools using Monte Carlo simulations. PloS one 8, e64287 (2013).

11. Z. A. Knight, M. E. Feldman, A. Balla, T. Balla, K. M. Shokat, A membrane capture assay for lipid kinase activity. Nature Protocols 2, 2459–2466 (2007).

12. L. P. Deanne et al., Mutations in DivL and CckA rescue a divJ null mutant of *Caulobacter crescentus* by reducing the activity of CtrA. Journal of Bacteriology 188, 2473–2482 (2006).

13. H. K. Zane, J. K. Doh, C. A. Enns, K. E. Beatty, Versatile Interacting Peptide (VIP) Tags for Labeling Proteins with Bright Chemical Reporters. Chembiochem 18, 470–474 (2017).

14. K. Lasker et al., Selective sequestration of signalling proteins in a membraneless organelle reinforces the spatial regulation of asymmetry in *Caulobacter crescentus*. Nature microbiology 5, 418–429 (2020).

15. P. Li et al., Phase transitions in the assembly of multivalent signalling proteins. Nature 483, 336–340 (2012).

16. S. F. Banani et al., Compositional Control of Phase-Separated Cellular Bodies. Cell 166, 651–663 (2016).

17. S. Boeynaems et al., Spontaneous driving forces give rise to protein-RNA condensates with coexisting phases and complex material properties. Proc Natl Acad Sci U S A 116, 7889–7898 (2019).

18. K. M. Ruff, S. Roberts, A. Chilkoti, R. V. Pappu, Advances in Understanding Stimulus-Responsive Phase Behavior of Intrinsically Disordered Protein Polymers. J Mol Biol 430, 4619–4635 (2018).

19. S. Saurabh, A. M. Perez, C. J. Comerci, L. Shapiro, W. E. Moerner, Super-resolution imaging of live bacteria cells using a genetically directed, highly photostable fluoromodule. Journal of the American Chemical Society 138, 10398–10401 (2016).

20. P. D. Dahlberg et al., Cryogenic single-molecule fluorescence annotations for electron tomography reveal in situ organization of key proteins in *Caulobacter*. Proceedings of the National Academy of Sciences, (2020).

21. K. M. Ruff, F. Dar, R. V. Pappu, Ligand effects on phase separation of multivalent macromolecules. Proc Natl Acad Sci U S A 118, (2021).

22. S. Kroschwald, S. Maharana, A. Simon, Hexanediol: a chemical probe to investigate the material properties of membrane-less compartments. Matters 3, e201702000010 (2017).

23. R. J. Wheeler et al., Small molecules for modulating protein driven liquid-liquid phase separation in treating neurodegenerative disease. bioRxiv, 721001 (2019).

24. H. Yaginuma et al., Diversity in ATP concentrations in a single bacterial cell population revealed by quantitative single-cell imaging. Scientific reports 4, 6522 (2014).

25. M. D. Spalding, S. T. Prigge, Lipoic acid metabolism in microbial pathogens. Microbiol Mol Biol Rev 74, 200–228 (2010).

26. N. Ausmees, J. R. Kuhn, C. Jacobs-Wagner, The bacterial cytoskeleton: an intermediate filament-like function in cell shape. Cell 115, 705–713 (2003).

27. N. Al-Husini, D. T. Tomares, O. Bitar, W. S. Childers, J. M. Schrader, α-Proteobacterial RNA degradosomes assemble liquid-liquid phase-separated RNP bodies. Molecular Cell 71, 1027–1039.e1014 (2018).

28. R. Düster, I. H. Kaltheuner, M. Schmitz, M. Geyer, 1,6-Hexanediol, commonly used to dissolve liquid-liquid phase separated condensates, directly impairs kinase and phosphatase activities. Journal of Biological Chemistry 296, (2021).

29. K. Heinrich et al., Molecular Basis and Ecological Relevance of *Caulobacter* Cell Filamentation in Freshwater Habitats. mBio 10, e01557–01519 (2019).

30. A. Klosin et al., Phase separation provides a mechanism to reduce noise in cells. Science (New York, N.Y.) 367, 464–468 (2020).

31. C. Gao et al., Intrinsic disorder in protein domains contributes to both organism complexity and clade-specific functions. Scientific Reports 11, 2985 (2021).

32. G. K. Pattanayak et al., Daily Cycles of Reversible Protein Condensation in Cyanobacteria. Cell Reports 32, 108032 (2020).

33. Y. Pu et al., ATP-Dependent Dynamic Protein Aggregation Regulates Bacterial Dormancy Depth Critical for Antibiotic Tolerance. Molecular Cell 73, 143–156.e144 (2019).

34. S. C. Weber, A. J. Spakowitz, J. A. Theriot, Nonthermal ATP-dependent fluctuations contribute to the in vivo motion of chromosomal loci. Proceedings of the National Academy of Sciences 109, 7338–7343 (2012).

35. Bradley R. Parry et al., The Bacterial cytoplasm has glass-like properties and is fluidized by metabolic activity. Cell 156, 183–194 (2014).

## References

1. M. Thanbichler, A. A. Iniesta, L. Shapiro, A comprehensive set of plasmids for vanillate- and xylose-inducible gene expression in Caulobacter crescentus. Nucleic Acids Research 35, e137 (2007).

2. S. Saurabh, A. M. Perez, C. J. Comerci, L. Shapiro, W. E. Moerner, Super-resolution imaging of live bacteria cells using a genetically directed, highly photostable fluoromodule. Journal of the American Chemical Society 138, 10398–10401 (2016).

3. J. W. Chin et al., Addition of p-azido-L-phenylalanine to the genetic code of Escherichia coli. Journal of the American Chemical Society 124, 9026 (2002).

4. A. M. Perez et al., A localized complex of two protein oligomers controls the orientation of cell polarity. mBio 8, 2238 (2017).

5. P. D. Dahlberg et al., Identification of PAmKate as a red photoactivatable fluorescent protein for cryogenic super-resolution imaging. Journal of the American Chemical Society 140, 12310–12313 (2018).

6. C. A. Bayas et al., Spatial organization and dynamics of RNase E and ribosomes in Caulobacter crescentus. Proc Natl Acad Sci U S A 115, E3712–E3721 (2018).

7. G. R. Bowman et al., Oligomerization and higher-order assembly contribute to sub-cellular localization of a bacterial scaffold. Molecular Microbiology 90, 776–795 (2013).

8. D. G. Gibson et al., Enzymatic assembly of DNA molecules up to several hundred kilobases. Nature methods 6, 343–345 (2009).

9. T. H. Mann, W. Seth Childers, J. A. Blair, M. R. Eckart, L. Shapiro, A cell cycle kinase with tandem sensory PAS domains integrates cell fate cues. Nature communications 7, 11454 (2016).

10. Z. A. Knight, M. E. Feldman, A. Balla, T. Balla, K. M. Shokat, A membrane capture assay for lipid kinase activity. Nature Protocols 2, 2459–2466 (2007).

11. A. Ertel, A. G. Marangoni, J. Marsh, F. R. Hallett, J. M. Wood, Mechanical properties of vesicles. I. Coordinated analysis of osmotic swelling and lysis. Biophysical Journal 64, 426–434 (1993).

12. J. B. Grimm et al., A general method to improve fluorophores for live-cell and single-molecule microscopy. Nature methods 12, 244–250 (2015).

13. S. Saurabh, M. Zhang, V. R. Mann, A. M. Costello, M. P. Bruchez, Kinetically Tunable Photostability of Fluorogen-Activating Peptide–Fluorogen Complexes. ChemPhysChem 16, 2974–2980 (2015).

14. I. J. Domian, K. C. Quon, L. Shapiro, Cell type-specific phosphorylation and proteolysis of a transcriptional regulator controls the G1-to-S transition in a bacterial cell cycle. Cell 90, 415–424 (1997).

15. R. T. Wheeler, L. Shapiro, Differential Localization of Two Histidine Kinases Controlling Bacterial Cell Differentiation. Molecular Cell 4, 683–694 (1999).

16. Y. Pu et al., ATP-Dependent Dynamic Protein Aggregation Regulates Bacterial Dormancy Depth Critical for Antibiotic Tolerance. Molecular Cell 73, 143–156.e144 (2019).

17. A. Ducret, E. M. Quardokus, Y. V. Brun, MicrobeJ, a tool for high throughput bacterial cell detection and quantitative analysis. Nature microbiology 1, 16077 (2016).

18. J. Schindelin, et al., Fiji: an open-source platform for biological-image analysis. Nature methods 9, 676–682 (2012).

19. S. K. Radhakrishnan, M. Thanbichler, P. H. Viollier, The dynamic interplay between a cell fate determinant and a lysozyme homolog drives the asymmetric division cycle of *Caulobacter crescentus*. Genes & development 22, 212–225 (2008).

20. B. L. Sprague, J. G. McNally, FRAP analysis of binding: proper and fitting. Trends in Cell Biology 15, 84–91 (2005).

21. A. Gahlmann et al., Quantitative multicolor subdiffraction imaging of bacterial protein ultrastructures in three dimensions. Nano Letters 13, 987–993 (2013).

22. C. Bayas, A. von Diezmann, A.-K. Gustavsson, W. E. Moerner, Easy-DHPSF 2.0: open-source software for three-dimensional localization and two-color registration of single molecules with nanoscale accuracy. Protocol Exchange 1, (2019).

23. G. J. Schütz, H. Schindler, T. Schmidt, Single-molecule microscopy on model membranes reveals anomalous diffusion. Biophysical Journal 73, 1073–1080 (1997).

24. T. Savin, P. S. Doyle, Static and Dynamic Errors in Particle Tracking Microrheology. Biophysical Journal 88, 623–638 (2005).

25. X. Michalet, Mean square displacement analysis of single-particle trajectories with localization error: Brownian motion in an isotropic medium. Physical review. E, Statistical, nonlinear, and soft matter physics 82, (2010).

26. L. Weimann et al., A quantitative comparison of single-dye tracking analysis tools using Monte Carlo simulations. PloS one 8, e64287 (2013).

27. J. Huff, The Airyscan detector from ZEISS: confocal imaging with improved signal-to-noise ratio and super-resolution. Nature methods 12, i–ii (2015).

28. P. D. Dahlberg et al., Cryogenic single-molecule fluorescence annotations for electron tomography reveal in situ organization of key proteins in *Caulobacter*. Proceedings of the National Academy of Sciences, (2020).

