## Supplementary Info for "ATP-responsive biomolecular condensates tune bacterial kinase signaling"

**Affiliations:**
[1]Department of Developmental Biology, Stanford University School of Medicine, Stanford, CA, USA

[2] Department of Chemistry, Stanford University, Stanford, CA, USA

[3] Department of Plant Biology, Carnegie Institution for Science, Stanford, CA, USA

Current address:

† Machine Learning and Imaging, Indigo Ag., 500 Rutherford Ave, Boston, MA, USA

‡ Intelligent Imaging Innovations (3i), 3575 Ringsby Court, Suite 102, Denver, CO, USA

**Notes**

**Acknowledgments:** The authors thank Dr. Rohit Pappu, Dr. Jared Schrader, Dr. Sean Crosson and Dr. Sunish Radhakrishnan for helpful discussions on experimental aspects of the manuscript. The authors also thank Drs. Harley McAdams, Richard Losick, Kotaro Fujii, Dante Ricci, Edward de Koning and Bo Gu for a careful reading of the manuscript. The authors thank all members of the Shapiro and Moerner labs for their constructive feedback throughout this work.

**Funding:** This work was supported by the National Institute of General Medical Sciences, the National Institutes of Health (R35-GM118071 to L.S.), and (R35-GM118067 to W.E.M.). L.S. is a Chan Zuckerberg Biohub Investigator.

**Author contributions:** This study was conceived by S.S. Experiments were designed by S.S., with inputs from W.E.M. and L.S. Strains for this study were engineered by S.S. and T.N.C. Single molecule imaging and data analyses were performed by C.B. and S.S., with inputs from W.E.M. Confocal imaging was performed by H.N.C. and S.S. Electron microscopy was performed by P.D.D. and S.S. The manuscript was written by S.S. and L.S., with inputs from W.E.M.

**Competing interests:** Authors declare no competing interests

**Data and materials availability:** All data is available in the main text or the supplementary materials. All data, code, and cell strains used in this work are available upon request.

**Supplementary Materials**

Supplementary Figures - Figs. S1-S4

Captions for Movies

Materials and Methods

References

Movies S1 through S5 (attachment)

**Figures – Supplementary text**

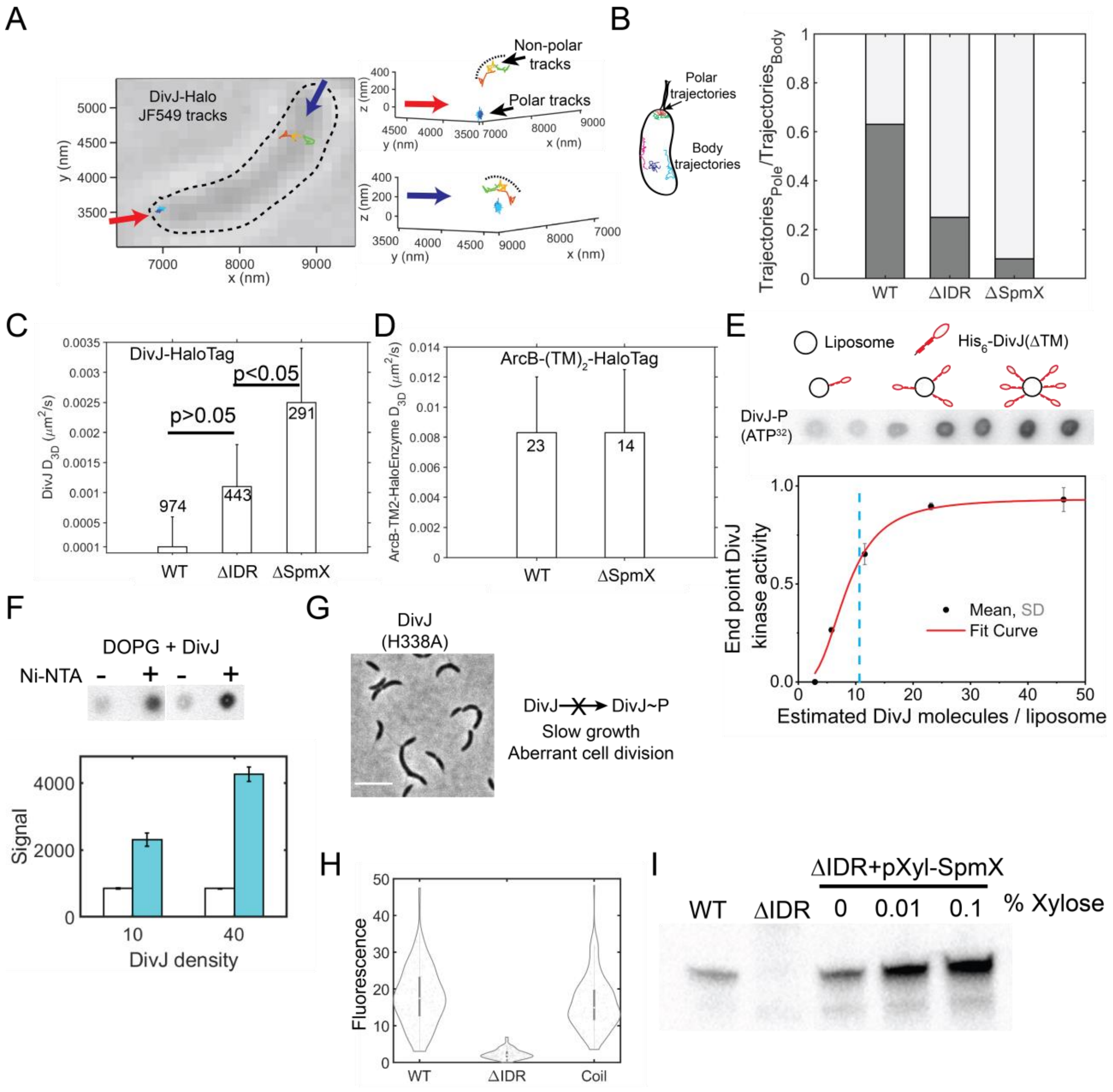


**Fig. S1. Effect of SpmX-IDR on DivJ localization, diffusion, and activity.** Throughout this panel, ΔIDR denotes SpmXΔIDR. **(A)** (left) Representative bright-field image of a live *Caulobacter* cell, overlaid with 2D projections of 3D DivJ-HaloTag trajectories measured using single-particle tracking. Each trajectory is plotted using a different color. Arrows denote perspectives for the 3D representation shown on the right. (right) Representation of the trajectories from the left panel in 3D space with appropriate perspective and all three Cartesian axes. Green and red trajectories in the body are membrane-associated as can be seen from their curvature, highlighted using a black dashed line. 3D tracking reveals polar as well as non-polar membrane associated trajectories. **(B)** Fraction of single molecule DivJ trajectories observed in the pole vs the body for respective strains. Total tracks analyzed are 1558 (WT), 1764 (ΔIDR) and 3572 (ΔSpmX). SpmX-IDR enables DivJ sequestration at the pole. **(C)** Three-dimensional diffusion coefficients obtained from Mean Squared Displacement (MSD) analyses for polar DivJ trajectories in various SpmX backgrounds. The number of trajectories analyzed for each sample are noted with the bars. P values from a pairwise two-tailed t-test are denoted in the plot. **(D)** 3D diffusion coefficients obtained from MSD analyses of control trans-membrane HaloTag trajectories in WT and SpmX deletion backgrounds. The HaloTag was attached to the membrane using two trans-membrane helices from the *E. coli* protein ArcB. Differences were not detected in the polar diffusion coefficients of the HaloTag in WT or SpmX deletion backgrounds. Errors in D in both panels (C) and (D) are standard deviations estimated from bootstrapping (50% sampling with replacement performed 500 times). **(E)** DivJ auto-kinase activity assayed on liposomes. Liposomes contained 90% (mol %) di-oleoyl-phosphatidyl glycerol (DOPG) and 10% (mol %) Nickel-chelated lipids (DGS-NTA) to bind $His_6$-DivJ(ΔTM). A fixed concentration of 5 μM $His_6$-DivJ(ΔTM) was pre-incubated with increasing amounts of liposomes to obtain increasing surface density of $His_6$-DivJ(ΔTM) molecules per liposome. Kinase reaction was performed in a buffer (25 mM HEPES-KOH, pH 7.4, 50 mM KCl, 25 mM NaCl, 5 mM $MgCl_2$) by adding 0.1 µCi ATP($\gamma$–$^{32}$P) for 5 min, quenching and blotting on a nitrocellulose membrane followed by phosphor imaging. The measured activity at each $His_6$-DivJ(ΔTM) surface density state was normalized to the highest density condition. Representative phosphor imaging data from spot assays and a schematic depicting increasing DivJ density (red) on liposomes (black circles) are shown above the graph. Intensity from phosphor imaging spots was quantified and the average data from triplicate spot assays were fit to a Hill-Langmuir curve as described in the methods section. Dashed blue line represents the average physiological polar DivJ concentration of 11 molecules measured from a single molecule counting assay for cells grown in M2G media. Error bars represent the standard deviation from 3 measurements. DivJ exhibited a density dependent kinase activity *in vitro*. **(F)** Liposome phosphorylation assay with (cyan bars) or without (white bars) Ni-NTA lipids at two different estimated densities of 10 and 40 DivJ molecules/ liposome. Error bars are standard deviations from 3 measurements. DivJ signal from liposomes without Ni-NTA lipids is insensitive to the protein density. **(G)** Representative phase contrast image of DivJ (H338A) cells in which

DivJ phosphorylation is absent, leading to aberrant cell division (scale bar 5µm). **(H)** Distribution of the ratio of mean polar and non-polar DivJ signal in WT cells (N=235), ∆IDR cells (N=258) and cells with SpmX∆IDR fused to a CoilZ peptide (N = 204) that can bind to a cognate CoilY peptide fused to DivJ-eYFP. Coil tag recruitment of DivJ gives a polar signal comparable to WT cells independent of the SpmX-IDR. **(I)** Western blot showing the levels of WT SpmX in WT, SpmX∆IDR, and SpmX∆IDR cells expressing WT SpmX on a chromosomal Xylose promoter (rescue cell line). For the rescue cell line, WT SpmX was produced by adding the indicated amount of Xylose. Western blot was performed using an antibody raised against the IDR of SpmX (*6*). Even without Xylose addition, the mild expression of SpmX was comparable to SpmX levels in WT cells.

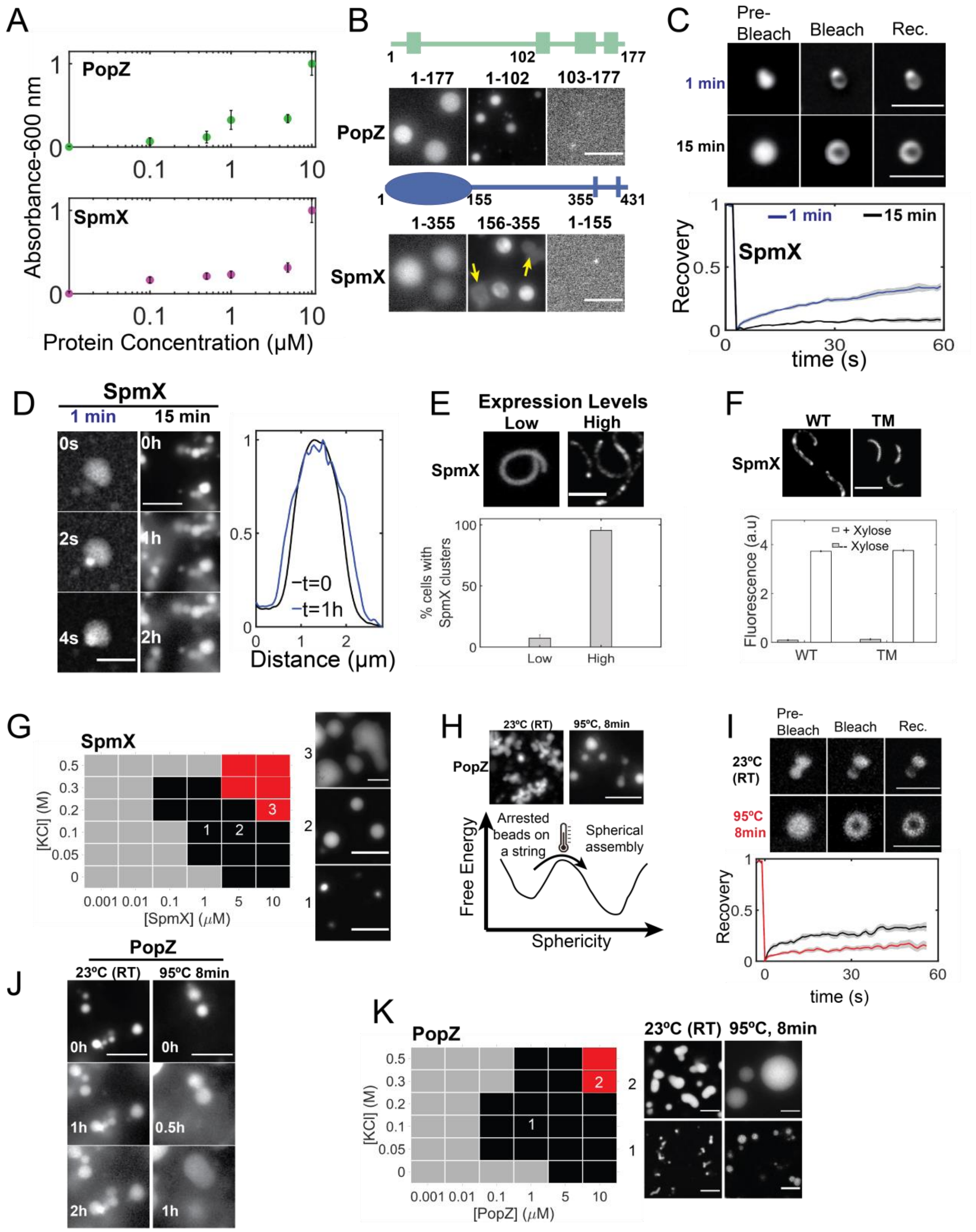


**Fig. S2. Context dependent properties of SpmX and PopZ condensates.** (All scale bars in this figure are 5 μm.) **(A)** Solution turbidity measured as a function of increasing PopZ (top) or

SpmX(ΔTM) (bottom) concentration in a physiological buffer (50 mM HEPES-KOH, pH 7.4, 0.1 M KCl) at room temperature. Background subtracted-absorbance values were normalized to the maximum absorbance observed in each case. Error bars represent standard deviation from 3 independent measurements. PopZ and SpmX exhibit a concentration dependent increase in turbidity. **(B)** (top) Representative fluorescent micrographs of PopZ domains: PopZ (full length), PopZ (AA 1-102) and PopZ (AA 103-177) (all proteins 1% Atto488 labeled). (bottom) Representative fluorescent micrographs of SpmX domains: SpmX (ΔTM, 5% Cy3 labeled), eYFP-tagged SpmX-IDR, and SpmXΔIDR (ΔTM, 5% Cy3 labeled). Yellow arrows in eYFP-tagged SpmX-IDR sample indicate condensates that degraded upon contacting the coverslip. All proteins are at 5 µM concentration. IDR containing regions of both SpmX and PopZ are necessary and sufficient for phase separation *in vitro*. **(C)** Internal rearrangement of SpmX condensates was assayed by FRAP at two different time points after fusing on glass. (top) A 250 nm spot was bleached in a SpmX condensate (5 µM protein, ΔTM, 5% Cy3 labeled), and the change in the fluorescence was analyzed as a function of time for condensates relaxed on glass within 1 min (blue) or after 15 mins (black). (bottom) Analyses of the fluorescence recovery of 1-min-old (black; N = 9), 15-min-old (green; N = 11) condensates are shown. Gray shadow depicts the standard error of the mean. SpmX condensates exhibit concentration dependent dynamics *in vitro*. **(D)** (Left) Condensates of SpmX fused to glass within 1 min (5 µM protein, ΔTM, 5% Cy3 labeled) displayed spontaneous fusion on the seconds time scale, and (Right) condensates older than 15 mins ripen on the hours' time scale without fusion. Measurement of the diameter of a ripening SpmX condensate over 1 hour showed an increase in its diameter by ~350 nm. Ripening is associated with slow condensate internal dynamics as a result of protein self-association. **(E)** Fluorescence micrographs of *Caulobacter* cells harboring a *popZ* deletion and over-expressing eYFP-labeled SpmX on a chromosomal Xylose promoter, induced using 0.03% Xylose (left) or 0.3% Xylose (right) for 1 hour. Bar graph below shows the % of cells exhibiting two or more SpmX clusters per cell (N ~ 400 cells for each case). SpmX cluster formation is concentration dependent *in vivo*. **(F)** Representative fluorescence micrographs of cells expressing SpmX-eYFP in a *popZ* deletion (left) or SpmX(TM)-eYFP (Δ1-355) in a *spmX* deletion, on a chromosomal Xylose promoter, induced using 0.3% Xylose for 1 hour. While clusters were observed for WT SpmX (>95% cells), SpmX-TM (Δ1-355) always remained diffuse. Shown below are end point YFP fluorescence measured using a plate reader in cells expressing SpmX-eYFP or SpmX-TM-eYFP (Δ1-355) grown in the presence or absence of 0.3% Xylose for 1 hour. At comparable concentrations, SpmX-TM domain without the cytoplasmic domain was unable to form clusters *in vivo*. **(G)** Phase diagram of SpmX as a function of protein and KCl concentration. Black squares denote conditions under which >20 fluorescent spots were observed to be above the intensity threshold (<background> + $3\sigma_{background}$) from 80 µm square regions across six fields of view spanning two biological replicates. Red squares denote metastable condensates suggestive of the spinodal decomposition on the phase diagram. Representative fluorescent micrographs from conditions 1-3 are shown next to the plot. **(H)** Representative maximum intensity projections of 3D confocal micrographs of PopZ (5 µM, 1% Atto 488 labeled) reconstituted in a physiological buffer (50 mM HEPES-KOH, pH 7.4, 0.1 M KCl). PopZ condensates exhibited "beads on a string" morphology at 23 ˚C (left) but relaxed into spherical condensates upon heat treatment (right). This suggests the presence of a thermal barrier between conformational states of different sphericity, depicted in the schematic below. **(I)** Internal rearrangement of PopZ condensates (5 µM, 1% Atto488 labeled) *in vitro* was assayed by FRAP. (top) A 250 nm spot was bleached, and the change in the fluorescence was analyzed as a function of time for untreated condensates (black) and annealed condensates (red). (bottom) Analysis of fluorescence recovery of untreated condensates (black; N = 10), annealed condensates (red; N = 12) are shown. Gray shadow depicts the standard error of the mean. Annealed condensates exhibit slower internal dynamics compared to arrested condensates. **(J)** Representative fluorescence micrographs showing ripening behavior of arrested PopZ condensates (5 µM protein, 1% Atto488 labeled) without fusion on the hours' time scale *in vitro*. Annealed PopZ condensates fused over sub-1 hour time scales showing the differences between the slow dynamic states of condensates *in vitro*. **(K)** Phase diagram of PopZ condensates (1% Atto488 labeled) as a function of protein and KCl concentration. Black squares denote conditions under which >20 fluorescent spots were observed to be above the intensity threshold (<background> + $3\sigma_{background}$) from 80 µm square regions across five fields of view spanning two biological replicates. Red squares denote metastable condensates suggestive of a spinodal decomposition on the phase diagram. Representative fluorescent micrographs from conditions 1-2 for the untreated and heat treated samples are shown next to the plot.

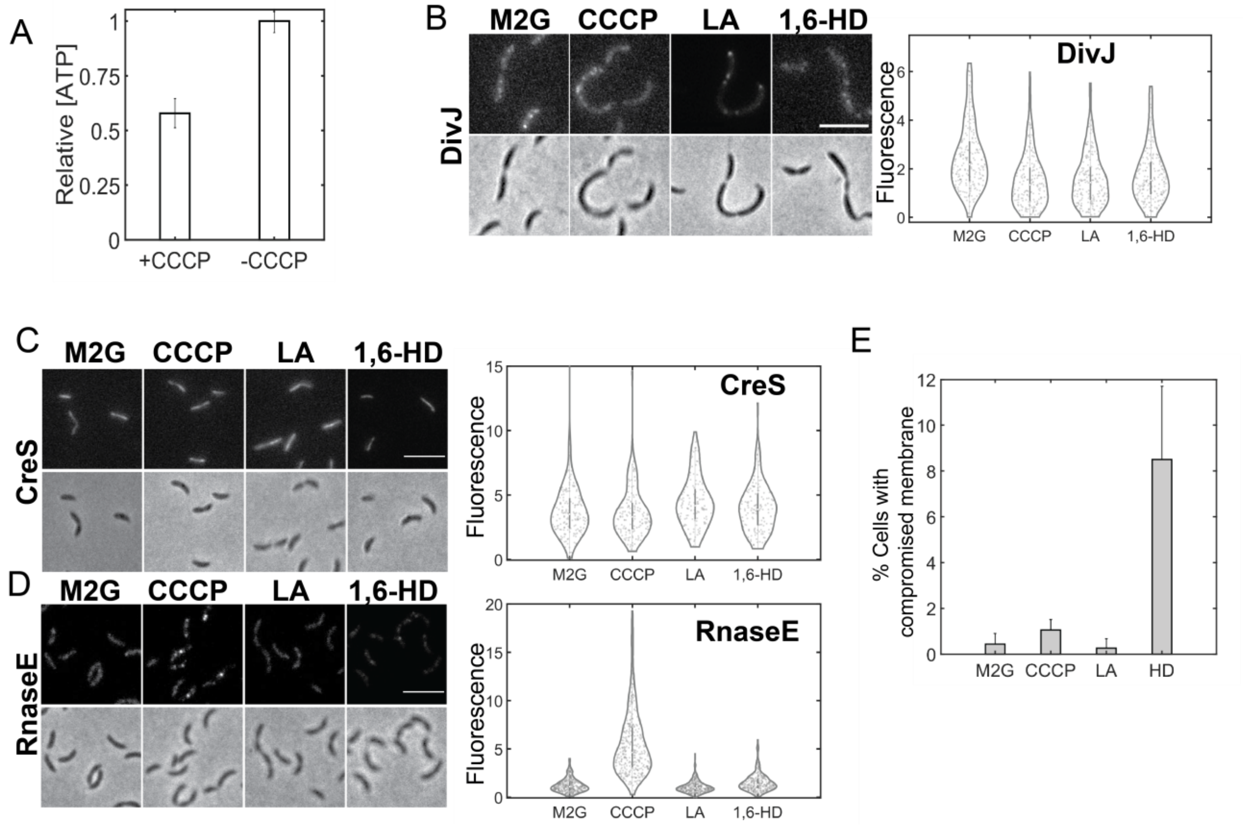


**Fig. S3. Effect of solutes on various protein assemblies and membrane integrity *in vivo*.** (All scale bars in this figure are 5 µm.) **(A)** Relative intracellular ATP concentrations in *Caulobacter* cells treated with 100 µM CCCP dissolved in DMSO, or an equivalent volume of DMSO for 10 minutes, measured using a commercial Luciferase based assay. Background from the cell growth media was subtracted and luminescence units were normalized to the ATP levels in the DMSO (-CCCP) case. Error bars represent the standard deviation from 3 biological replicates, with each sample assayed in triplicate. CCCP addition leads to rapid ATP depletion in live *Caulobacter* cells. **(B)-(D)** Effect of ATP depletion by CCCP addition (100 µM, 10 min), 5 µM LA, or 5% (v/v) 1,6-HD treatment (30 mins, each) on various protein assemblies *in vivo*. **(B)** Representative fluorescence (top) and phase contrast (bottom) micrographs of *Caulobacter* cells with *spmX* deletion and over-expressing DivJ-eYFP on a high copy Xylose inducible plasmid exhibited fluorescent clusters. However, the fluorescence within DivJ-eYFP clusters was depleted under all treatments (CCCP, LA, 1,6-HD). ~600 cells were analyzed for each condition. **(C)** Representative fluorescence and phase contrast micrographs of *Caulobacter* cells over-expressing Cres-eYFP exhibited fluorescent fibers. The relative fluorescence intensity in these fibers depleted under CCCP treatment, while a mild enhancement in fluorescence was observed under HD or LA treatment. ~500 cells were analyzed for each condition. **(D)** *Caulobacter* cells expressing endogenously tagged RnaseE-eYFP exhibited clusters with enhanced fluorescence under CCCP treatment (10 mins) and depleted fluorescence under LA treatment but not under HD treatment (30 mins, each). ~800 cells were analyzed for each condition. ATP and LA, but not HD, are able to dissolve condensates such as BR bodies while having different effects on structured protein assemblies such as CreS, or protein aggregates such as DivJ in the absence of SpmX *in vivo*. **(E)** Bar plot showing percentage of *Caulobacter* cells expressing eYFP-PopZ with membrane leakage observed under treatment with CCCP (N = 762 cells), LA (N=682 cells) and HD (N = 474 cells), compared with the untreated cells (M2G, DMSO, N = 719 cells). Cells were treated with the respective solutes followed by labeling with Live-or-Dye$^{TM}$ (640/662) dye based on manufacturer's protocols. Dye labeled cells were washed thrice using M2G and imaged on an epifluorescence microscope. Cells with compromised membranes exhibited at least five-fold higher fluorescence compared to healthy cells. This threshold criterion was applied to all the images to obtain the bar plots. Error bars are standard deviation across 12 fields of view from three biological replicates. 1,6-HD treated cells exhibited 4-5-fold higher membrane leakage than CCCP or LA treated cells.

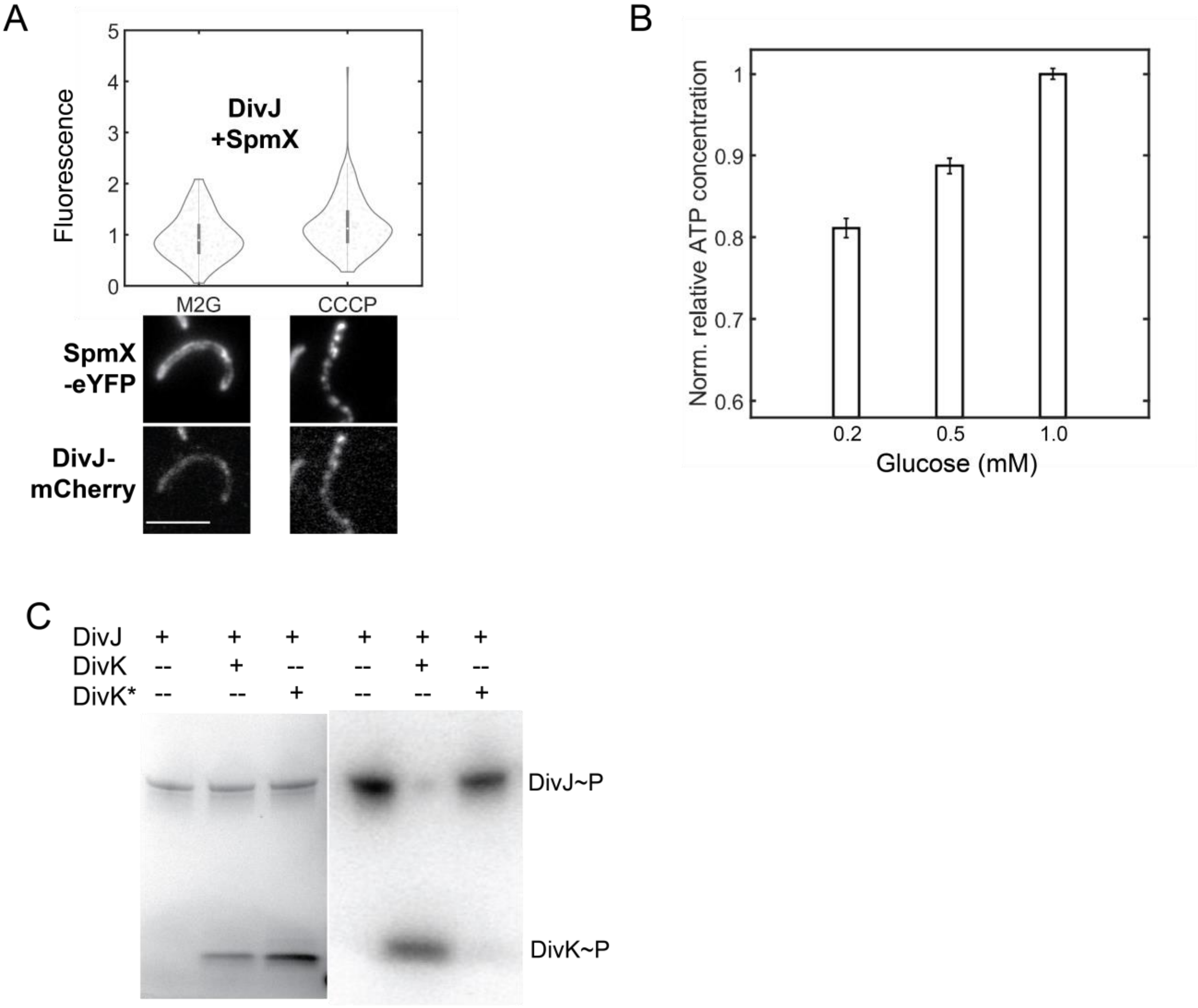


**Fig. S4. Effect of ATP on DivJ localization and phenotype. (A)** Distribution of the ratio of polar localized to diffuse DivJ signal in *Caulobacter* cells co-expressing SpmX-eYFP and DivJ-mCherry in a strain harboring *popZ* deletion under the control of Xylose and Vanillate promoters, respectively. DivJ-mCherry was expressed using 50 mM Vanillate while SpmX-eYFP cluster formation was induced by the addition of 0.3% Xylose for 4 hours. Cells were treated with DMSO (M2G) or 100 µM CCCP dissolved in DMSO (10 mins) followed by imaging. Representative fluorescence micrographs of cells in both eYFP and mCherry channels are shown below (scale bar is 5 µm). ~400 cells were analyzed for each condition. Polar localization of DivJ is enhanced under ATP depletion (scale bar 5µm). **(B)** Bar plot showing the relative intracellular ATP concentrations in *Caulobacter* cells grown in M2 minimal media as a function of glucose concentration from 0.2 mM to 1 mM. ATP concentrations were measured using a commercial Luciferase based luminescence assay. Error bars represent standard error of the mean from 3 biological replicates each assayed in triplicate. **(C)** *In vitro* phosphorylation measurement of the cytoplasmic domain of DivJ (5 μM) alone, in the presence of DivK (5 μM), or DivK* (DivK(D53N)) (5 μM) incubated with 1 μCi ATP($\gamma$–$^{32}$P) in a physiological buffer (50 mM HEPES-KOH, pH 7.4, 0.1 M KCl, 5 mM $MgCl_2$) for 5 mins. Phosphorylation reactions were quenched by addition of SDS, and proteins were separated via SDS-PAGE. (left) Silver-stained SDS-PAGE gel with DivJ, DivJ and DivK or DivJ and DivK* (Right) Radiograph showing the phosphorylation levels of the respective proteins from the gel on the left. The cytoplasmic domain of DivJ is capable of autophosphorylation and phosphor-transfer to DivK, but not a mutant of DivK that lacks its phospho-acceptor amino acid residue (D53).

**Movie Captions**

**Movie S1**

Three dimensional widefield fluorescence images of condensates formed by eYFP fusion of the SpmX-IDR. 5 µM eYFP-tagged SpmX-IDR was incubated in a physiological buffer (50 mM HEPES-KOH, pH 7.4, 0.1 M KCl) for 30 mins followed by microscopy. Slices from 0-0.8 µm show the presence of collapsed eYFP-IDR condensates on glass surface treated with aminosilane. Slices above 1 µm reveal the presence of spherical condensates diffusing in the buffer.

**Movie S2**

Sequential two-color, three dimensional, widefield fluorescence imaging of condensates formed by eYFP-SpmX-IDR and DivJ. 5 µM eYFP-tagged SpmX-IDR was incubated in the presence of 5 µM DivJ (ΔTM, 1% labeled with JF646) in a physiological buffer (50 mM HEPES-KOH, pH 7.4, 0.1 M KCl) for 30 mins followed by microscopy. The SpmX and DivJ channels are separated in time by 8 ms. DivJ is enriched within SpmX-IDR condensates *in vitro*.

**Movie S3.**

3 dimensional confocal imaging of PopZ condensates at room temperature. 5 µM PopZ (1% labeled with Atto488) was reconstituted in a physiological buffer (50mM HEPES-KOH, pH 7.4, 0.1 M KCl) in an aminosilane treated glass bottom chamber for 30 mins. PopZ exhibited an arrested, beads on a string-like morphology that formed an extended network of condensates in 3 dimensions. The 3D stack was acquired using a 63x oil objective with 280 nm slices.

**Movie S4.**

3 dimensional images of PopZ were acquired as in Movie S3 followed by deconvolution in Fiji. A 12 µM by 12 µM by 6 µM volume was selected from a Z-stack and exported into Imaris. A Gaussian filter spanning 1 pixel was applied to the image and 3D reconstruction was performed. Various perspectives of the condensates were then exported into a movie.

**Movie S5.**

Observation of dynamic hollow condensates of 5 µM SpmX (ΔTM, 5% labeled with Cy3) incubated in the presence of 1 mM ATP in a physiological buffer (50 mM HEPES-KOH, pH 7.4, 0.1 M KCl, 5 mM $MgCl_2$) at room temperature for 30 minutes. Images were captured every 10 seconds and the movie was recorded for 25 minutes. The jitter in the hollow condensates is due to bubble fusion events along the z-axis (into the plane of the image).

### Materials and Methods

#### 1. Strain Engineering

| **Table S1. Plasmids and strains used in the study** | | |
|---|---|---|
| **A. Plasmids** | | |
| pXYFPC-2 | pxyl:eYFP *Caulobacter* integrating plasmid | (*1*) |
| pXYFPC-4 | pxyl:eYFP *Caulobacter* integrating plasmid | (*1*) |
| pVCHYC-2 | pvan:CHY Caulobacter integrating plasmid | *(1)* |
| pMCS-4 | *Caulobacter* integrating plasmid | (*1*) |
| pMCS-6 | *Caulobacter* integrating plasmid | (*1*) |
| pNPTS-138 | *Caulobacter* counter selection plasmid | (*1*) |
| pAP510 | pspmX:spmX-dL5 | (*2*) |
| pAP515 | pdivJ:divJ-dL5 | (*2*) |
| pSS083 | pdivJ:divJ-HaloTag | This work |
| pSS225 | pdivJ:divJ-HaloTag-CoilY | This work |
| pTS35 | spmX(1-162)-eYFP-CoilZ-spmX(356-431) | This work |
| pTS18 | pdivJ:divJ-eYFP | This work |
| pTS23 | spmX(1-162)-eYFP-spmX(356-431) | This work |
| pEvol-pAzF | Plasmid to incorporate p-Azido-L-phenylalanine | Addgene (*3*) |
| pSS47 | pet28a-popZ-TEV-His6 | This work |
| pSS121 | pet28a-spmX(1-355)-F338AzF-TEV-His10 | This work |
| AP434 | pTev5-divJ(D1-187)-His6 | (*4*) |
| pSS102 | pet28a-popZ(1-102)TEV-His6 | This work |
| pSS103 | pet28a-popZ(103-177)TEV-His6 | This work |

| | | |
|---|---|---|
| pSS120 | pet28a-spmX(1-155)-TEV-His10 | This work |
| pSS165 | pet28b-spmX(Δ1-155)eYFP-(156-355)-TEV-His10 | This work |
| pAP549 | pXyl:spmX | (*4*) |
| pAP519 | pBX:spmX-eyfp | (*4*) |
| pTC276 | pNTS138-upstream divK-divK-Halo-3xFlag-downstream divK | This work |
| pTC8 | pBX-mCherry-popZ | This work |
| pAP565 | pspmX:spmx(1-155)-dL5 | (*4*) |
| pSS306 | pV:divJ-mCherry | This work |
| pSS194 | pspmx:spmX-PAmKate | (*5*) |
| pSS319 | pBX-DivJ-eYFP | This work |
| ***B. Bacterial*** **strains** | | |
| AP414 | pspmX:spmX-eYFP | (*2*) |
| AP369 | ΔspmX | (4) |
| AP510 | pspmX:spmX-dL5 | (*2*) |
| SS087 | pdivJ:divJ-HaloTag | This work |
| TS4 | pdivJ:divJ-HaloTag, pspmX:spmX-dL5 | This work |
| TS5 | ΔspmX, pdivJ:divJ-HaloTag | This work |
| TNC17 | pBX-mCherry-PopZ | This work |
| TS15 | pdivJ:divJ-eYFP | This work |
| SS297 | ΔpopZ pBXyl-SpmX-eYFP | This work |
| TS34 | ΔspmX pspmX:spmX(1-162)-eYFP-spmX(356-431) | This work |

| | | |
|---|---|---|
| SS057 | pet28a-PopZ-TEV-His6 | This work |
| SS150 | pXyl-ArcbTM2-HaloTag | This work |
| SS159 | ΔSpmX-pXyl-ArcBTM2-HaloTag | This work |
| AP451 | pXyl-CreS-eYFP | (*2*) |
| JS51 | pranseE:RnaseE:eYFP | (*6*) |
| SS311 | ΔpopZ pBXyl-SpmX-eYFP pVDivJ-mCherry | This work |
| SS235 | ΔspmX pspmX:spmX(1-162)-eYFP-CoilZ-spmX(356-431) ΔdivJ pdivJ:divJ-HaloTag-CoilY | This work |
| SS324 | ΔspmX pBX-DivJ-eYFP | This work |
| SS294 | ΔspmX pspmX:spmX(1-162)-eYFP-spmX(356-431), pdivK:divK-HaloTag-3xFlag | This work |
| SS295 | ΔspmX pspmX:spmX-dL5, pdivK:divK-HaloTag-3xFlag | This work |
| ***C. Primers*** | | |
| TC29F | gagttttggggagacgaccatatgGTGAGCAAGGGCGAGGAGGATAAC | |
| TC30R | ggttcttgagactgatcggacatGGTACCATGCATATTAATTAAGGCGCC | |
| TC532F | CTTCGTCGTAATTGCCGGGATTGG | |
| TC533R | TCGTTGTCGTCGACGATCAGCACC | |
| TC540F | agctacgtaatacgactcactagtGGGTCGACAGGTCGGCCAGGGC | |
| TC541R | ttaaggtaccTGCAGGCTGCCTTTCCAGCAGG | |
| TC544F | acgatgacaagGCATGAGCGCCCGGATCCTCG | |
| TC545R | gtcacggccgaagctagcgaattcCTTGACGCGCGCGGACAGTTCC | |
| SS013F | ctttaagaaggagatatacatggctatgtccgatcagtctca | |

| | |
|---|---|
| SS014R | ggagctcgaattcggatctcagtggtggtggtggtggtgctccgtGCTCTGAAAATACAGGTTTTCgtaggcgccgcgtccccga |
| SS7F | tctagaaataattttgtttaactttaagaaggagatatacATGAAACCGCGTCATCAGGT |
| SS8R | cgacggagctcgaattcgtcagtgatggtggtggtggtgatggtgatggtgGCTCTGAAAATACAGGTTTTCCTCCACCAGCGGCACGTC |
| SS015F | ctttaagaaggagatatacatggctatgtccgatcagtctca |
| SS016R | gagctcgaattcggatctcagtggtggtggtggtggtgctccgtGCTCT |
| SS017F | actttaagaaggagatatacgacgaagtcgccgagcagctggtcggcgtt |
| SS018R | gcaagcttgtcgacggagctcgaattcggatctcagtggtggtggtggtggtgctccgtGCTCT |
| SS9F | tctagaaataattttgtttaactttaagaaggagatatacATGAAACCGCGTCATCAGGT |
| SS10R | cgacggagctcgaattcgtcagtgatggtggtggtggtgatggtgatggtgGCTCTGAAAATACAGGTTTTCCCATTCGCCGTTGGCCGG |
| SS321F | aaccacgatgcgaggaaacgcatatgTtgGAATTCGAAACGCTTCC |
| SS322R | cggagctcgagatcttaaggtaccGCGCGGCGCAAAGGCGATGACG |

| **Table S2. Filters used** | | |
|---|---|---|
| Fluorophore | Filter (Excitation \| Dichroic \| Emission) | Supplier |
| CFP | 436/20 \| 455 \| 480/40 | Semrock |
| Atto 488 | 480/40 \| 505 \| 527/30 | Semrock |
| eYFP | 500/20 \| 515 \| 535/30 | Semrock |
| mCherry/ Cy3/ JF549 | 554/23 \| 573 \| 609/54 | Semrock |
| dL5/ Cy5/ JF646 | 635/18 \| 652 \| 680/42 | Semrock |

$$\mathrm{MSD}(\tau) = 2dD\left(\tau - \frac{\Delta t}{3}\right) + 2\sum_{j=1}^{d} {\sigma_j}^2 \qquad \text{Equation 2}$$

In Equation 2 above, $\tau$ is time lag, $\Delta t$ is camera exposure time, $d$ is the number of dimensions (3 in this case), $D$ is the diffusion coefficient, and $\sigma_j$ is the localization error in dimension $j$.
